# Engineering a light-responsive, orthogonal Lon protease in *E. coli* for targeted protein degradation

**DOI:** 10.64898/2026.06.17.732960

**Authors:** Cristian Coriano-Ortiz, Lily A. Fenton, Josh Kome, Mary J. Dunlop

## Abstract

Optogenetic methods are powerful tools for synthetic biology, allowing light to control cellular processes. While most bacterial optogenetic systems regulate gene expression at the transcriptional level, relatively few enable post-translational control, which can provide faster and growth-independent regulation of protein activity. Here, we describe the development of a post-translational optogenetic tool in Escherichia coli using the Mesoplasma florum Lon (mfLon) protease, an AAA+ protease that is orthogonal to native E. coli degradation machinery. To engineer a light-responsive mfLon, we constructed a large library in which a blue-light responsive domain, Avena sativa LOV2, was introduced into nearly every codon position in the protease using an unbiased molecular approach. We screened 726 mfLon-LOV variants using fluorescence-activated cell sorting and multi-round enrichment campaigns. We identified a novel dark-active variant (mfLon-LOV-534) that degrades target proteins in the dark and is inactivated upon blue-light exposure. Characterization of this variant demonstrates that its proteolytic response can be tuned by varying blue-light intensity and transcriptional expression levels. Furthermore, we show that mfLon-LOV-534 can degrade a target protein in both exponential and stationary growth phases, which addresses the limitations of division-based protein dilution. This work establishes a scalable approach to engineering allosteric control in complex multimeric enzymes and provides a foundation for orthogonal, growth-independent control of protein stability in synthetic circuits.

## Introduction

Optogenetic systems are an attractive choice for controlling synthetic circuits because they provide precise, tunable, and reversible control over cellular processes.^1–5^ In bacterial optogenetics, the emphasis has been on transcriptional control, enabling applications probing regulatory networks,^6,7^ controlling metabolic flux,^8,9^ and precisely regulating gene expression.^1,3,4,10,11^ While these applications have demonstrated the clear utility of light in controlling cellular processes, genetic regulatory networks consist of more than transcriptional regulators. Many cellular processes adapt to changing environmental conditions by directly altering enzymatic function. For example, metabolic networks can achieve this adaptation through the allosteric control of enzymes. Optogenetic tools that directly control protein function offer the potential to better mimic these systems.

Bacterial optogenetic systems that directly control protein function exist, but often still focus on transcriptional regulation. For example, the popular optogenetic systems CcaS-CcaR^12^ and OptoCre^13^ both allosterically control enzyme function, but the ultimate outcome is regulation of gene expression. A downside of this transcriptional focus is that the majority of proteins in *E. coli* are not actively degraded; protein clearance occurs through cell division, thereby limiting the dilution rate to the organism’s doubling time.^14^ During stationary phase or slow-growth conditions, proteins persist, and make transcriptional repression ineffective. Natural regulatory systems mitigate this limitation via post-translational control with targeted degradation mechanisms. Post-translational tools can decouple cellular processes from gene expression delays and growth dependence by modulating control at the protein level. Recently, optogenetics research has begun to explore active protein degradation as a means of control.^15–18^

The systems opto-TEVp^16^, LOVdeg^15^, and OptoGPlad^18^ are recent optogenetic tools for targeted protein degradation. All three work by regulating the exposure of a degron on the protein of interest, though the specific mechanism varies: opto-TEVp controls the activity of a site-specific protease that exposes a cryptic degron upon light excitation, LOVdeg is translationally fused to a target protein and exposes a sequestered degron after light excitation through a conformational change, and OptoGPlad adds a phosphate group to an arginine of the target protein leading to its degradation. All three systems rely on *E. coli*’s endogenous proteases to degrade the target protein, but none of them directly control the protease’s enzymatic activity. Endogenous *E. coli* proteases, such as *E. coli* Lon, play a key role in orchestrating the degradation of misfolded proteins and regulatory proteins. Therefore, direct optogenetic control of these native proteases is likely to be detrimental to maintaining vital physiological processes in the cell.

Engineering a light-responsive protease orthogonal to native *E. coli* machinery is an attractive option, as it shifts the point of control from the current strategy of exposing a degron to the protease itself while still addressing the need for post-translational control. Allosteric control of an orthogonal protease, as opposed to induction of its expression, also offers the potential to bypass the delay of *de novo* protease synthesis, potentially allowing for rapid activation of the degradation pathway. Controlling protein degradation via a protease would also address the limitations of division-based protein clearance by providing an “off-switch” that can clear proteins in a manner that is independent of cell growth. Further, using orthogonal degradation pathways reduce the risk of interfering with endogenous processes. This mechanism can also serve as a modular approach to mimic allosteric control of enzymes via targeted degradation.

In this work, we engineered a novel optogenetic protein degradation system by allosterically controlling the enzymatic activity of *Mesoplasma florum* Lon protease (mfLon). mfLon is a hexameric AAA+ protease that can be expressed in *E. coli* but acts on an ssrA tag that is orthogonal to *E. coli*’s native degradation machinery.^19^ To render mfLon light-responsive, we used a variant of *Avena sativa* LOV2 (AsLOV2), a popular domain for optogenetic allosteric control. AsLOV2 is widely used for engineering optogenetic proteins due to several beneficial features, including its small size, defined conformational change, and proximity of the N- and C-termini in its caged state.^20,21^ AsLOV2 has a C-terminal Jα helix docked to the protein’s core when kept in the dark. Upon exposure to blue light (∼450 nm), the Jα helix undocks, and the region becomes unstructured. This conformational change has been exploited to impact protein function when inserted within a target enzyme.^20–22^ However, engineering allosteric control remains a challenging endeavor due to its context dependence and the lack of clear rules governing which insertions yield functional, switchable proteins.

Historically, the approach to engineer allostery using AsLOV2 and other domains has been through rational protein design.^21,23,24^ Rational identification of potential domain insertion sites typically requires a 3D structure of the target protein to identify tightly packed, surface-exposed loops near functional domains. At the same time, highly conserved residues should be avoided, as evolutionary conservation suggests they may be essential for protein folding or function. Lastly, potential sites should be mapped onto the protein’s 3D structure to ensure they accommodate the structural perturbation of domain insertion.^21^ While rational design following these guidelines can be powerful, studies have shown that functional insertion sites can also exist outside of these described search criteria^20,25^ and that meeting all of these criteria does not guarantee achieving the desired function.^23^ In addition, engineering allosteric control of mfLon poses three additional challenges. First, mfLon is a homohexamer; thus, the insertion of a light-responsive domain must coordinate conformational changes across all mfLon protomers to avoid unintended disruption of the enzymatic activity. At present, few studies exist on the rational design of multimers.^21^ Second, each protomer of mfLon is relatively large at 787 amino acids, meaning there are many possible domain insertion sites. Third, there is no crystal structure of mfLon, which is essential for rational design; thus, its domain classifications are based solely on sequence homology.

We initially attempted to use rational design rules to identify insertion sites within mfLon, but these studies failed to produce a functional variant. Next, we used a random, unbiased approach to target every site within mfLon. We built a domain-insertion library with AsLOV2 by using the saturated programmable insertion engineering (SPINE)^26^ protocol. We screened the library for functional variants by employing an enrichment campaign using fluorescence-activated cell sorting (FACS). Sorted populations were sequenced, and we identified potential functional variants via enrichment analysis. We compared the results of our enrichment analysis with predicted functional sites from rational protein design and with the ProDomino^27^ machine learning model. Remarkably, after screening 726 variants, we identified a single insertion site that yielded a functional light-responsive mfLon protease. These results highlight the protease’s sensitivity to domain insertions and the need for high-throughput screening of complex multimeric targets.

Here, we present data characterizing mfLon-LOV-534 as a novel post-translational optogenetic tool for targeted protein degradation in *E. coli*. We demonstrate that its proteolytic activity can be modulated by varying the intensity of blue light and the concentration of arabinose used for induction. Critically, mfLon-LOV-534 enables active protein clearance in both exponential and stationary growth phases, with rates faster than natural protein decay and independent of division-based dilution. By shifting control to the degradation machinery itself via an orthogonal protease, this tool provides a lever for post-translational regulation in synthetic circuits and biological engineering applications where precise, rapid, and growth-independent protein-level control is essential.

## Results

### Predicted mfLon structure and its function

To engineer light-controlled allosteric regulation, we sought to find functional insertion sites for AsLOV2 within the mfLon protease. The AsLOV2 domain comprises residues 404-546 of *Avena sativa* phototropin 1; previous reports have shown that truncating the domain can lead to tighter coupling of conformational change.^9,25^ Native AsLOV2 exhibits spontaneous undocking of the Jα helix that can occur in the absence of light. Because mfLon is a hexamer, an AsLOV2 insertion would be present in every protomer; therefore, we wanted to limit the potential for spontaneous conformational changes to maintain all AsLOV2 domains in a consistent state. To address this concern, we truncated the C-terminal end of the domain by three amino acids for tighter conformational coupling.^9,28^ We also introduced mutations L493V, H519R, V520L, D522G, G528A, E537F, N538Q, and D540A, which have been shown to result in tighter dark caging of the Jα helix.^15,29^ All mutations relative to the original AsLOV2 sequence are summarized in Fig. S1. From here on out, we refer to this modified AsLOV2 variant as ‘LOV’ for simplicity and to insertion chimeras of mfLon and LOV as ‘mfLon-LOV.’

Our goal was to identify functional insertional sites that propagate the light-induced conformational change in the LOV domain to mfLon, thereby disrupting enzymatic activity. To measure proteolytic activity, we created a reporter plasmid with an IPTG-inducible promoter driving expression of mChartreuse^30^ fused to a 27 amino acid mfLon degron, pdt.^31^ This mChartreuse-pdt design enabled us to couple protease activity to mChartreuse fluorescence readout (Fig. 1A). In the dark, we expected to see active degradation of the fluorescent reporter. Under blue light, our design goal was to disrupt proteolytic activity, resulting in fluorescent protein accumulation. Before attempting to insert LOV into mfLon, we confirmed that the protease itself was not light-responsive by inducing mfLon expression with arabinose and measuring fluorescence of mChartreuse-pdt under dark and blue light conditions (Fig. 1B). In the absence of arabinose induction, there were low levels of mfLon expression so mChartreuse levels remained high, regardless of light exposure. With 1mM arabinose, mfLon was expressed, leading to degradation of mChartreuse. In both cases, mfLon showed little response to blue light, confirming that the original protease was not light responsive.

**Figure 1.**
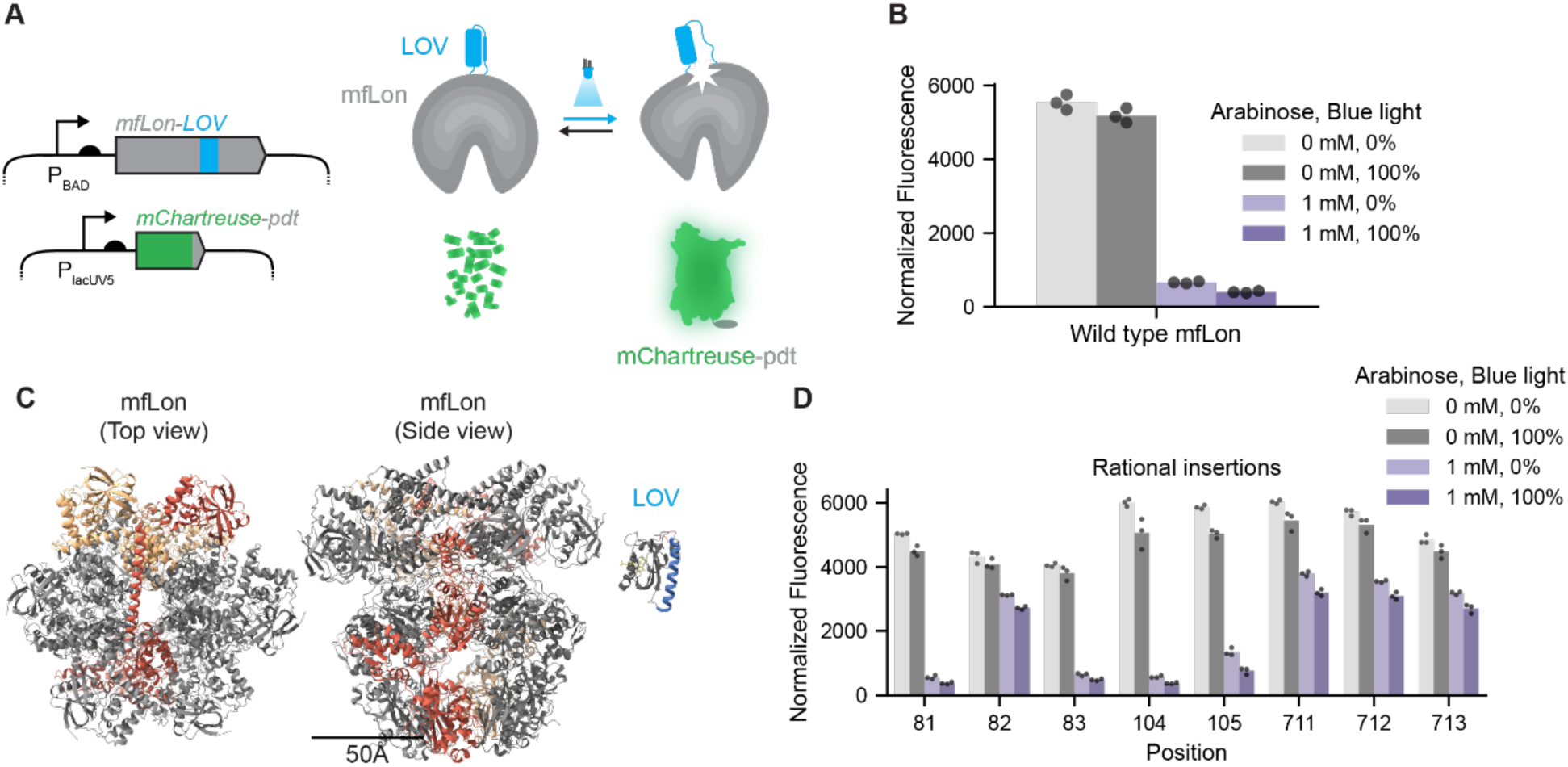
Rational design of light-responsive mfLon-LOV. (A) Schematic showing the expected mechanism of an mfLon-LOV variant with dark-induced degradation, where LOV’s conformational change disrupts proteolytic activity. Gene circuit design schematics show that arabinose induces mfLon-LOV expression from the P_BAD_ promoter. mChartreuse-pdt is induced with IPTG from the P_lacUV5_ promoter. (B) Wild-type mfLon degrades mChartreuse-pdt and is not light responsive. (C) Predicted structure of mfLon generated with AlphaFold3. Red and orange regions highlight two protomers of the hexameric structure. (D) Functional assay for rationally designed mfLon-LOV variants measuring mChartreuse-pdt fluorescence under different arabinose and light conditions. Normalized fluorescence corresponds to raw fluorescence divided by optical density at 600 nm. mChartreuse-pdt expression is induced with IPTG (100 µM). Blue light, 460 nm, at 100% corresponds to 1 mW/cm^2^. n = 3 technical replicates.

We first sought to find a functional insertion site using a rational design approach, following the methodology outlined by Dagliyan et al.^21^ Briefly, rational insertion sites are typically identified by searching for residues in tightly packed, surface-exposed loops that are not evolutionarily conserved and fall near a catalytically active area. Since mfLon lacks an experimentally determined structure, we used AlphaFold3 to predict its hexameric structure (Fig. 1C).^32^ Each protomer of mfLon is 787 amino acids, which is relatively large compared to LOV’s 140 amino acids. The model captures mfLon’s barrel shape, constructed by interacting protomers that are radially distributed, forming a cavity running through the middle of the structure through which a target protein is unfolded and degraded. Of note, the N-terminal domains of the protomers pair with those on opposite sides of the hexamer; this has been described in the literature as a trimer of dimers and is captured in the model (Fig. 1C).^33,34^ Using the predicted structure and the Pfam protein database, we identified that the N-terminal region functions as the degron recognition site, the middle region contains a highly conserved ATPase domain, and the C-terminal domain serves as the proteolytic site (Fig. S2).^35^ The model also shares key features with the cryo-EM structure of *Thermus thermophilus* Lon (ttLon, PDB: 7P6U) (Fig. S3), capturing important qualitative features such as the paired globular N-terminal domains situated above the ATPase domain.^34^ To further verify structural accuracy, we aligned the model against additional Lon homologs from *Escherichia coli* (PDB: 6U5Z), *Meiothermus taiwanensis* (PDB: 4YPL), and *Yersinia pestis* (PDB: 6ON2) and found consistent domain organization (Fig. S4). We note that the N-terminal domain of Lon is disordered, making it challenging to image and therefore absent from these structures.^36^

With a working model of mfLon’s structure, we applied the rational design criteria and identified three surface-exposed loops: residues 81-83, 104-105, and 711-713. Residues 81-83 and 104-105 reside in the N-terminal domain, while residues 711-713 reside in the C-terminal domain (Fig. S5-7). We constructed and tested variants at all proposed rational insertion sites, yielding a total of eight designs. In each case, our naming convention places the LOV insertion immediately after the numbered residue. For example, mfLon-LOV-81 has LOV placed between residues 81 and 82 of mfLon. Among the eight rational variants, we found no functional designs (Fig. 1D, Fig. S8). Sites 81, 83, 104, and 105 were inert, i.e., had no effect on mfLon’s enzymatic activity. Insertion at residues 82, 711, 712, and 713 exhibited a nonfunctional phenotype, i.e., almost complete loss of mfLon’s proteolytic activity. Although we failed to find a functional design among the rational variants, the results hinted at interesting effects depending upon minor changes in the insertion location. For example, we observed a significant change in protease behavior when the insertion site was shifted by just one amino acid from residue 81 to 82, and again from 82 to 83. These results suggest that modest differences in placement can have a dramatic impact on protease function. This sensitivity, combined with our unsuccessful search for rational insertion sites, motivated our shift to comprehensive library screening. This approach allowed us to broaden our search beyond the constraints of rational design and reduced the risk of overlooking a functional insertion site.

### Library generation and screening

We used the SPINE method to generate an mfLon domain-insertion library.^26^ The mfLon sequence is 2361 bp, corresponding to 787 amino acids, per protomer. Using SPINE’s Python-based pipeline, we divided the mfLon sequence into 13 fragments of approximately 60 amino acids each (Fig. 2A). For each fragment, we designed an oligo subpool comprised of ∼60 individual sequences flanked by external BsmBI cut sites and encoding an internal BsaI cut site after every codon. Simultaneously, for each of the 13 fragments, we designed primer pairs to amplify the plasmid containing mfLon, excluding the ∼60-amino-acid sequence of that fragment. After amplification, we inserted the oligo subpool into the backbone, replacing the omitted subsection via Golden Gate assembly (Fig. 2B). We repeated this process for every fragment, yielding an intermediate library containing BsaI cut sites after every codon in mfLon. To select for successful insertion of internal BsaI sites, we next introduced a chloramphenicol cassette flanked by BsaI cut sites with internal BsmBI cut sites into the intermediate library via Golden Gate assembly, yielding the second intermediate library (Fig. 2C). Throughout the construction process, we handled each fragment separately to facilitate troubleshooting and identify any regions that proved difficult to clone. This approach revealed that the subpool for fragment 13 (positions 728–787) was not successfully inserted into the backbone, whereas the other 12 fragments were assembled without issue. Because functional domain insertions are less likely in this region due to a high concentration of conserved residues, we proceeded with fragments 1-12, corresponding to comprehensive domain insertions between positions 1 and 727. To ensure proper coverage, we monitored colony-forming units (CFU) for each fragment. We confirmed that the number of transformants exceeded the number of possible variants in the sublibrary by an order of magnitude (>600 CFUs per fragment).

**Figure 2.**
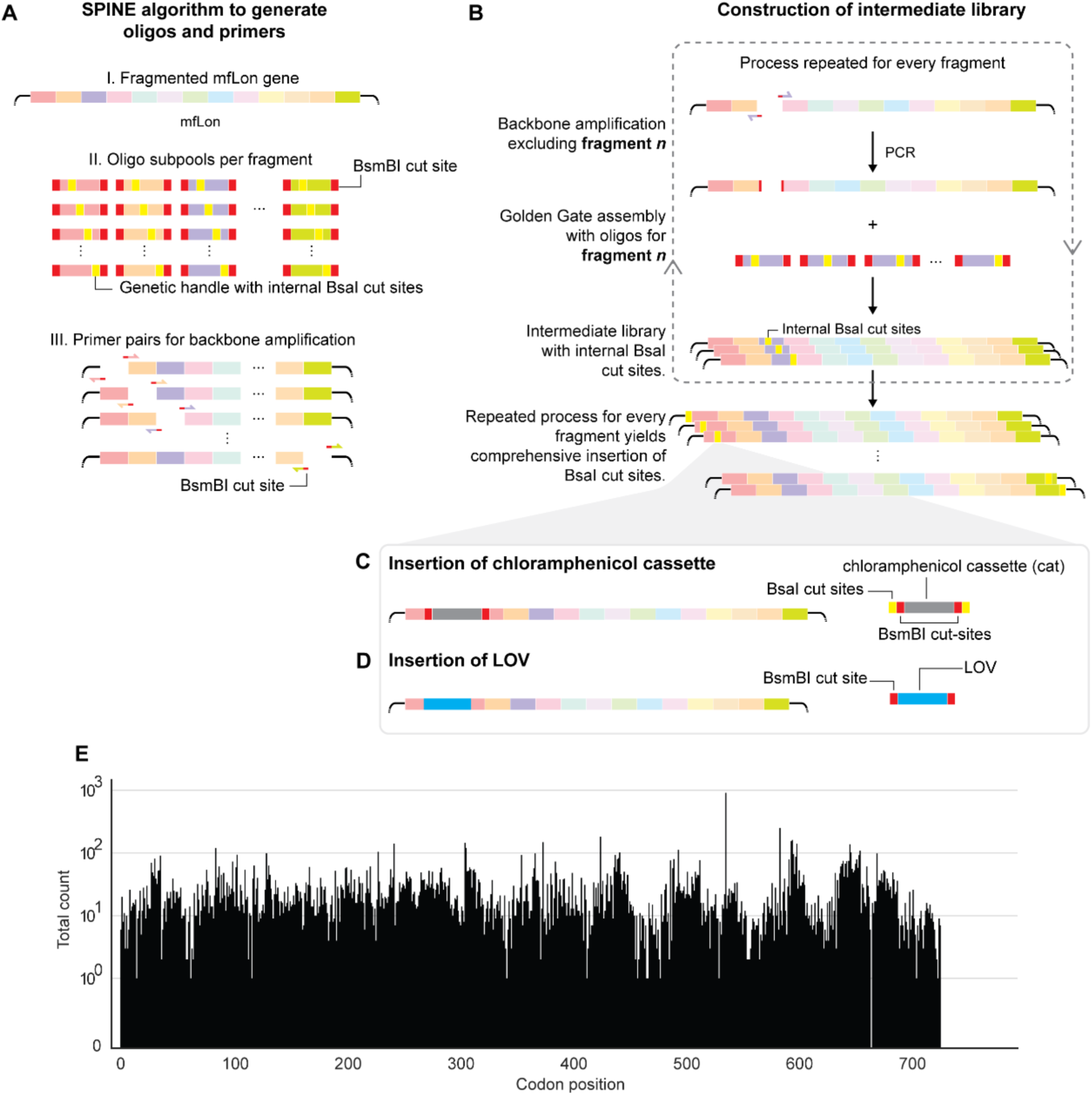
mfLon-LOV library generated using SPINE has >90% coverage. (A) Using the SPINE Python pipeline, we segmented the mfLon sequence into 13 fragments and generated oligo subpools for each fragment containing internal BsaI cut sites. We also generated primers for backbone amplification. (B) SPINE workflow to generate intermediate library. Plasmid containing mfLon is amplified to exclude fragment. The oligo subpool is inserted to generate variants via Golden Gate assembly. The process is repeated for each of the 13 fragments. (C) Chloramphenicol cassette is introduced via Golden Gate using the internal BsaI sites on the intermediate library. (D) LOV domain is inserted via Golden Gate using BsmBI cut sites, generating mfLon-LOV library. (E) Sequencing coverage of LOV insertion at mfLon codon position. The denoted codon position is the residue that immediately precedes the LOV insertion site.

To assemble the finalized library, we digested the BsmBI cut sites within the chloramphenicol cassette and inserted the LOV domain via Golden Gate assembly (Fig. 2D). We sequenced the resulting mfLon-LOV library, finding that we achieved nearly complete coverage (92.2%) of all possible insertion sites (Fig. 2E). The exceptions were insertions after codons 666 and 728-787, which accounted for just 7.8% of sites. Overall, using SPINE, we generated a nearly comprehensive library of insertion sites for LOV within mfLon, with relatively uniform coverage across the coding sequence.

### Sorting to find functional variants

To screen the library for functional variants, we used FACS to sort the library under different light conditions. To simplify the experimental workflow during sorting, we replaced the IPTG-inducible promoter in our initial reporter plasmid with a constitutively active promoter driving mChartreuse-pdt expression. We co-transformed cells with the mChartreuse-pdt reporter and the mfLon-LOV library. Functional mfLon-LOV variants that allosterically control protease activity could, in principle, take on one of two forms (Fig. 3A): (i) Dark-deg: The previously discussed case with dark-induced degradation. (ii) Light-deg: Light-induced degradation, with the inverse behavior. We reasoned that we would be more likely to isolate dark-deg variants than light-deg variants, as Jα helix undocking is more likely to disrupt than restore protein function. Indeed, this format, where undocking leads to protein disruption, is commonly used in the engineering of optogenetic proteins with LOV insertions.^20–23^ Nevertheless, there are isolated examples of Jα helix undocking restoring function,^20,37^ so we designed our sorting assay to include this possibility.

**Figure 3.**
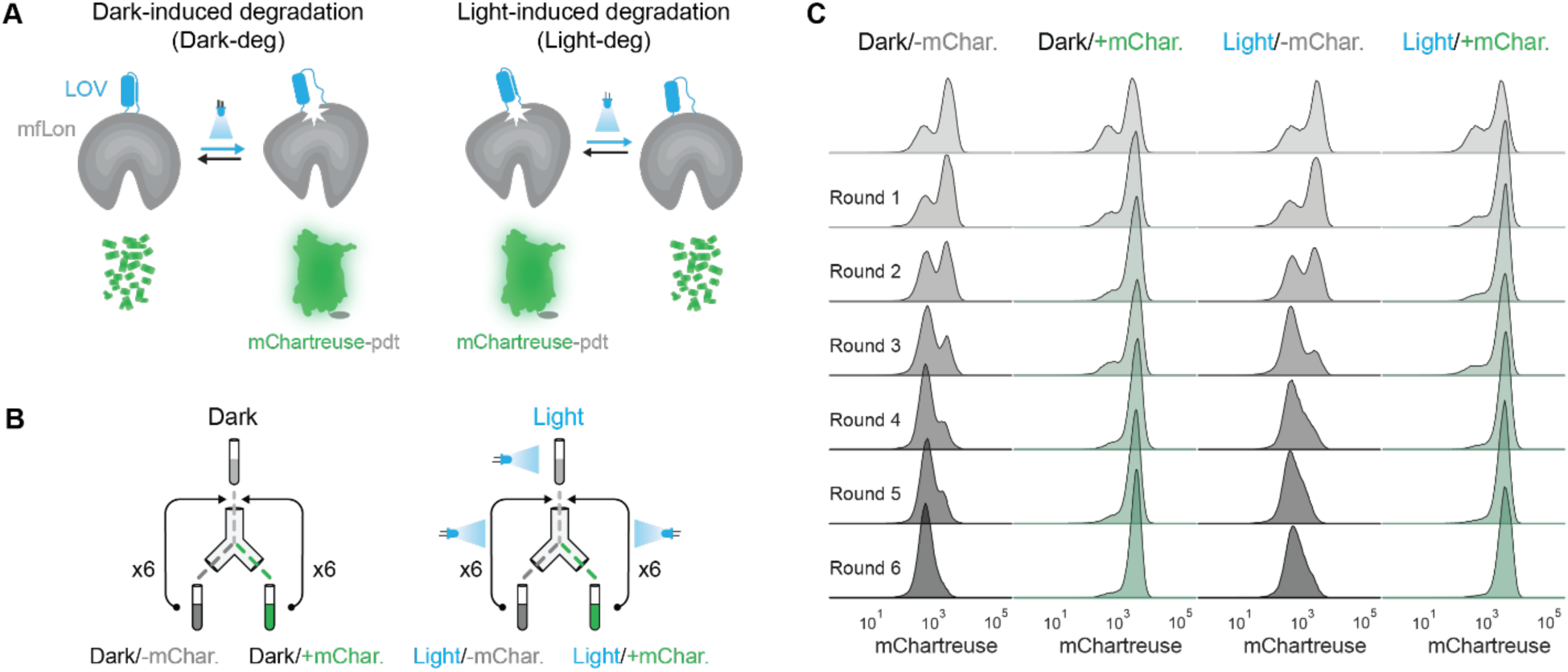
mfLon-LOV functional assay and enrichment. (A) Schematics representing possible functional variants. Dark-deg: Dark-induced degradation variants exhibit proteolytic activity in the dark, whereas light exposure disrupts degradation. Light-deg: Light-induced degradation variants exhibit impaired proteolytic activity in the dark, while light exposure restores activity. (B) Schematic representing FACS enrichment campaign. Cells co-transformed with the mfLon-LOV library and the mChartreuse-pdt reporter were kept in the dark or exposed to blue light, then sorted by fluorescence phenotype, grown for 16 hours, and subsequently returned to the same light conditions and gated as before. The process was repeated for six rounds. (C) Fluorescence distribution for mfLon-LOV library per bin condition and enrichment round. Every distribution shows 200,000 events.

In addition to the dark-deg and light-deg forms we hoped to isolate, there were two additional functional classes we expected to see: (iii) Inert: In these variants, the site of LOV insertion does not significantly impair protease activity, so fluorescent read-outs are low regardless of light exposure conditions. (iv) Nonfunctional: These LOV insertions significantly impair protease activity, so fluorescence read-outs are high regardless of light exposure conditions. Based on our initial findings with the rationally designed library variants (Fig. 1D), we expected the majority of variants to be either inert or nonfunctional in response to light, representing either benign but ineffective insertions or catastrophic insertion impacts.

Using the mChartreuse-pdt reporter with the mfLon-LOV library, we performed multiple rounds of enrichment with FACS (Fig. 3B). To set gates for sorting, we used a control strain co-transformed with the reporter plasmid and a plasmid containing an arabinose-inducible mfLon (Fig. S9). Using mChartreuse levels from the uninduced and induced control strain, we defined gates for high (+mChartreuse) or low (-mChartreuse) fluorescence. Cells containing the mfLon-LOV library were kept either in the dark or exposed to blue light. These populations were then sorted using the −/+mChartreuse gates, resulting in four bins: Dark/-mChartreuse, Dark/+mChartreuse, Light/-mChartreuse, and Light/+mChartreuse (Fig. 3B). These bins were subsequently enriched by growing the sorted populations under the same light conditions and sorting them again using the same fluorescence gates for six rounds. The initial fluorescence distributions of the library under both light conditions were bimodal, with a bias toward higher fluorescence. This indicates that most insertions impaired mfLon’s ability to degrade mChartreuse-pdt (Fig. 3C). Subsequent rounds of enrichment showed a progressive shift from bimodal to unimodal distributions corresponding to the desired fluorescence state. This shift suggests that we successfully isolated cells with the specified fluorescence properties.

### Establishing a framework for variant characterization

After the multi-round enrichment process, we sequenced the four resulting populations. In addition, we sequenced the naïve library present before sorting to serve as a basis for comparison of enriched sequence variants. We filtered for reads containing LOV insertions and counted the number of times each insertion site appeared in the naïve library and the four enrichment bins. In all cases, we obtained >40,000 reads per bin, which is >50x more than the number of possible variants, providing confidence that our sequencing had sufficient coverage.

We assessed enrichment and depletion of variants with different insertion sites by adapting the DIP-seq pipeline to calculate enrichment scores.^38^ Briefly, raw read counts were normalized by the total reads in each bin to yield relative frequencies, which were then divided by the naïve library frequency and log2-transformed. Each insertion site has a resulting value, called an enrichment score, that can be positive or negative. For positive values, the insertion site is enriched, i.e., it indicates an overrepresentation of a variant relative to the naïve library. For negative values, the insertion site is depleted.

Both FACS and sequencing are subject to experimental noise. We reasoned that relying on absolute enrichment scores could be misleading if a variant appears enriched in multiple bins due to sorting “leakiness” or sequencing bias. For example, 5.1% of insertion sites had a positive enrichment score exclusively in the Light/+mChartreuse bin and negative scores in all other bins, a pattern that is not biologically coherent and likely reflects noise (Fig. S10). To mitigate this and simplify variant characterization, we defined two metrics: ΔDark (Dark/-mChartreuse minus Dark/+mChartreuse enrichment score) and ΔLight (Light/-mChartreuse minus Light/+mChartreuse enrichment score). This approach serves as an internal normalization. For example, if a variant is enriched in both Dark/-mChartreuse and Dark/+mChartreuse bins relative to the naïve library, a positive ΔDark value indicates that the variant was more abundant in the active protease bin (Dark/-mChartreuse). In general, a positive Δ value indicates enrichment in the –mChartreuse bin (low fluorescence, active protease) relative to the +mChartreuse bin; a negative value indicates the opposite.

By plotting variants in ΔDark-ΔLight space, we categorized insertions based on their location with each quadrant belonging to a functional category. Variants with positive ΔDark and positive ΔLight values are expected to be inert, negative ΔDark and positive ΔLight are possible light-deg candidates, negative ΔDark and negative ΔLight are likely nonfunctional, and positive ΔDark and negative ΔLight are possible dark-deg variants (Fig. 4A). The ΔDark-ΔLight framework categorized 64% of variants as nonfunctional, 27% as inert, 3% as light-deg, 1.6% as dark-deg, and the remainder were not classifiable since they fell right on the x- or y-axis of the ΔDark-ΔLight space.

**Figures 4.**
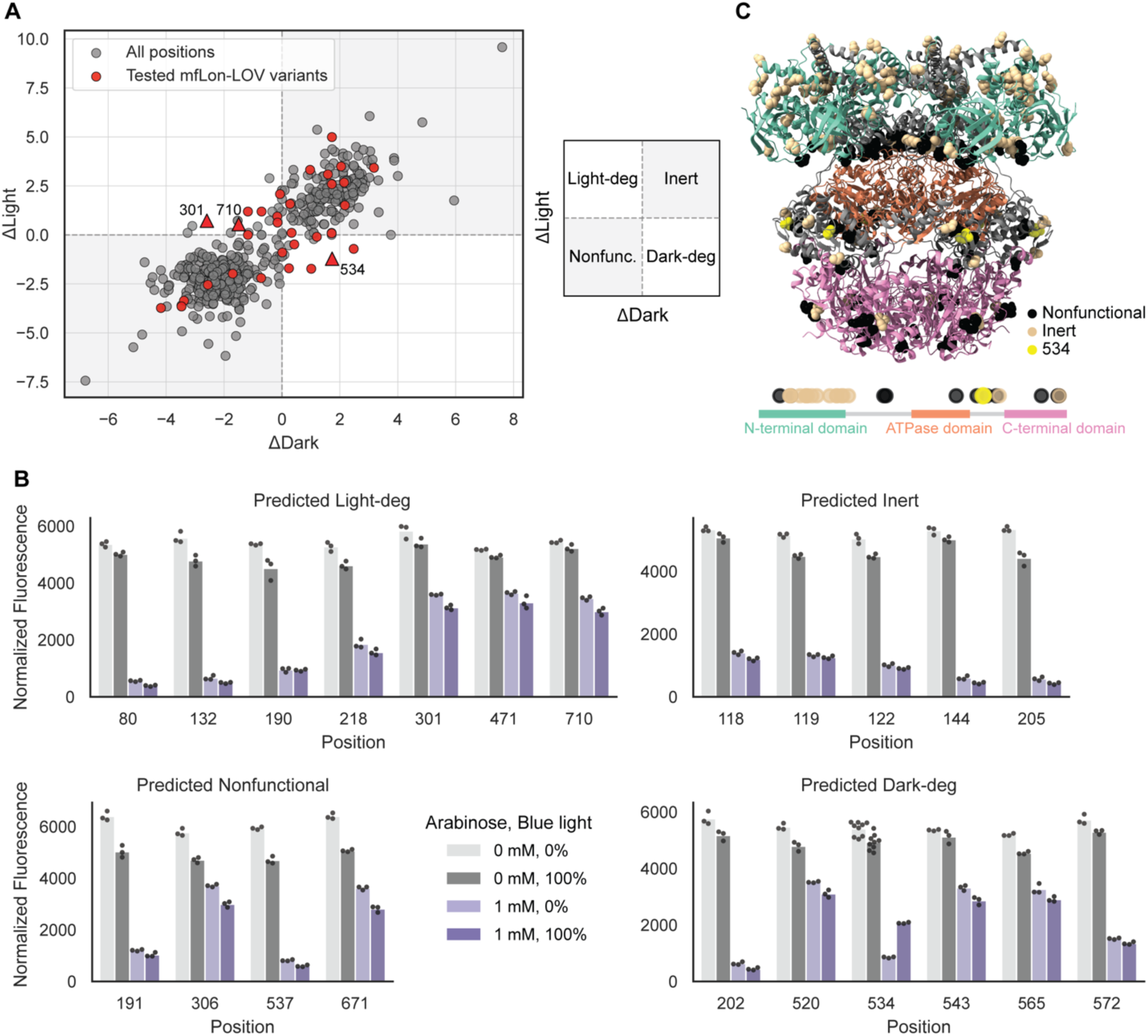
Testing library variants for functionality based on enrichment patterns. (A) Visualizing enrichment scores of variants in the ΔDark-ΔLight space. Quadrants can be used to predict functional categorization of mfLon-LOV variants. In total, we tested 22 variants in addition to the 8 rational design variants, all of which are highlighted in red. The primary candidates (534, 301, 710) are denoted with triangles. Data from all sequenced variants are shown in gray. (B) Functional assay of mfLon-LOV variants. The results are divided into the predicted classes: light-deg, dark-deg, inert, or nonfunctional. Normalized fluorescence corresponds to raw fluorescence divided by optical density at 600 nm. n ≥ 3 biological replicates. (C) Predicted mfLon structure with annotated locations of insertion sites based on their actual—not predicted—function. In addition, the locations of the sites within the mfLon sequence are shown.

Within the light-deg and dark-deg quadrants of the ΔDark-ΔLight space, three of the candidates displayed unambiguous enrichment patterns even before internal normalization: a dark-deg candidate at insertion site 534 and two light-deg candidates at insertion sites 301 and 710. Their raw enrichment scores across all four bins suggested clear light-responsive behavior, indicating a more conservative criterion for functionality (i.e., they fell into the correct bins in Fig. S10). Specifically, site 534 exhibited positive, negative, negative, and positive enrichment scores in the Dark/−mChartreuse, Dark/+mChartreuse, Light/−mChartreuse, and Light/+mChartreuse bins, respectively. Sites 301 and 710 displayed the inverse pattern: negative, positive, positive, and negative. Only these three insertion sites satisfied this strict criterion, representing approximately 0.5% of possible variants (Fig. S10).

### Experimental validation and structural insights

We asked whether predictions resulting from the ΔDark-ΔLight space were accurate. To test this, we cloned 22 variants spanning all four quadrants of ΔDark-ΔLight space. This set included the three primary candidates (534, 301, and 710), 15 additional sites selected to represent the full range of predicted functional categories, and 4 candidates predicted by the machine learning model ProDomino^27^ (Table S1). The ProDomino predictions should be interpreted as variants that are expected not to be nonfunctional, i.e., could be light-deg, inert, or dark-deg. We co-transformed each mfLon-LOV variant with the IPTG-inducible mChartreuse-pdt reporter to quantify protease activity (Fig. 4B, Fig. S11).

Of the three primary candidates, we confirmed that insertion site 534 was light-responsive, displaying dark-deg functionality. Despite their strong signatures in the enrichment analysis, sites 301 and 710 were nonfunctional and exhibited impaired proteolytic activity and no detectable light responsiveness. Among the 15 additional sites we tested, experimental results were consistent with several predictions from the ΔDark-ΔLight framework: all 5 inert-predicted sites (119, 205, and the 3 ProDomino candidates 118, 122, and 144) were correctly categorized, and 2 of the 4 nonfunctional predictions (306 and 671) were confirmed. Of the rational insertion sites previously identified, insertions 81, 83, and 104–105 were correctly predicted to be inert, and insertions 712–713 were correctly predicted to be nonfunctional. Site 82 was incorrectly predicted as inert because it was nonfunctional in practice. Site 711 could not be classified, as its ΔLight component value of 0 placed it directly on the x-axis. However, none of the predicted dark-deg or light-deg sites outside of 534 display functional light-responsive proteolysis, making 534 the only experimentally validated light-responsive variant in this tested set. ProDomino candidate 80, which fell in the light-deg quadrant, also showed no light responsiveness. We had hoped that the ΔDark-ΔLight framework’s internal normalization would allow us to parse the inherent noise of sequencing. It appears to be good at categorizing inert variants but is prone to false positives for functional site predictions. Adding to the complexity of identifying functional sites is their rarity: even with more conservative criteria, we discovered only one functional site.

Residue 534 lies adjacent to the ATPase domain of mfLon, leading us to hypothesize that the LOV conformational change propagates to this domain and disrupts proteolytic activity (Fig. 4C). This is a surprising result, as site 534 does not satisfy many criteria for selection as an insertion site based on rational design rules. Although it is sufficiently surface-exposed (Fig. S6), it is relatively conserved (Fig. S5), suggesting that it might be a poor choice, and, further, it does not reside in a tightly packed region (Fig. S7). Together, these metrics indicate that this site would likely have been overlooked without the comprehensive library screen. Inert insertions were concentrated primarily in the N-terminal domain of mfLon, which is known to be intrinsically disordered (Fig. 4C). This distribution is consistent with the expectation that insertions in disordered regions are less likely to perturb the protein’s structural integrity or enzymatic function. Nonfunctional insertions were distributed throughout the protein with no discernible regional bias, indicating that disruption of mfLon activity can arise from perturbations across many structural contexts. The classifications in Fig. 4C correspond to the actual, experimentally-measured categories, not predictions.

### Characterizing, optimizing, and testing mfLon-LOV-534

Our comprehensive library screening identified a single light-responsive candidate, mfLon-LOV-534, that shows a 2.4x fold change between dark and light conditions. We first asked what mChartreuse fluorescence we could expect as the maximum. This is different from the 0 mM arabinose condition because it accurately accounts for expression of the protease. By co-transforming mfLon-LOV-534 with an mChartreuse reporter lacking the pdt tag, we observed lower fluorescence than in the 0 mM arabinose case. mfLon-LOV-534 reaches 60% of the maximum achievable fluorescence when exposed to blue light (Fig. S12).

We were interested in testing mfLon-LOV-534 further to assess its tunability in response to light or to chemical induction. To ensure saturation of LOV activation and to minimize effects on cell growth, we first tested light intensity, applying blue light between 0 and 1 mW/cm^2^ (0 to 100%). We found that the protease exhibits a linear response to light intensity (Fig. 4A, Fig. S13). Because mfLon-LOV-534 is arabinose-inducible, we also tested different concentrations of this chemical inducer and found that the dynamic range of the response can be tuned by changing induction concentrations (Fig. 4B). The fold change between dark and light conditions plateaued at concentrations above 1 mM arabinose.

We were also interested in asking whether it was possible to increase the fold change and focused on the design of the linkers flanking the LOV insertion as candidate sequences for optimization. During library construction, we inserted the LOV domain flanked by glycine-serine (GS) linkers into mfLon. We included this linker in our original design because prior reports indicate low success with naïve LOV domain insertion, whereas insertion using GS linkers has been effective.^27,39^ We hypothesized that changing the linker could improve the propagation of light-induced conformational changes from LOV to mfLon, thereby better disrupting proteolytic activity in the light. To test this, we swapped the GS linker in mfLon-LOV-534 for GSG, GPG, GP, G, no linker, and no linker with a 3-amino-acid truncation at the N-terminus of the LOV domain. Surprisingly, all linker swaps resulted in LOV insertions that were inert (Fig. 5C). This indicates that mfLon-LOV-534 is highly sensitive to linker design, with single amino acid changes abolishing light responsiveness. Furthermore, both longer and shorter linker designs disrupt function, indicating that the original GS design represents a local optimum.

**Figure 5.**
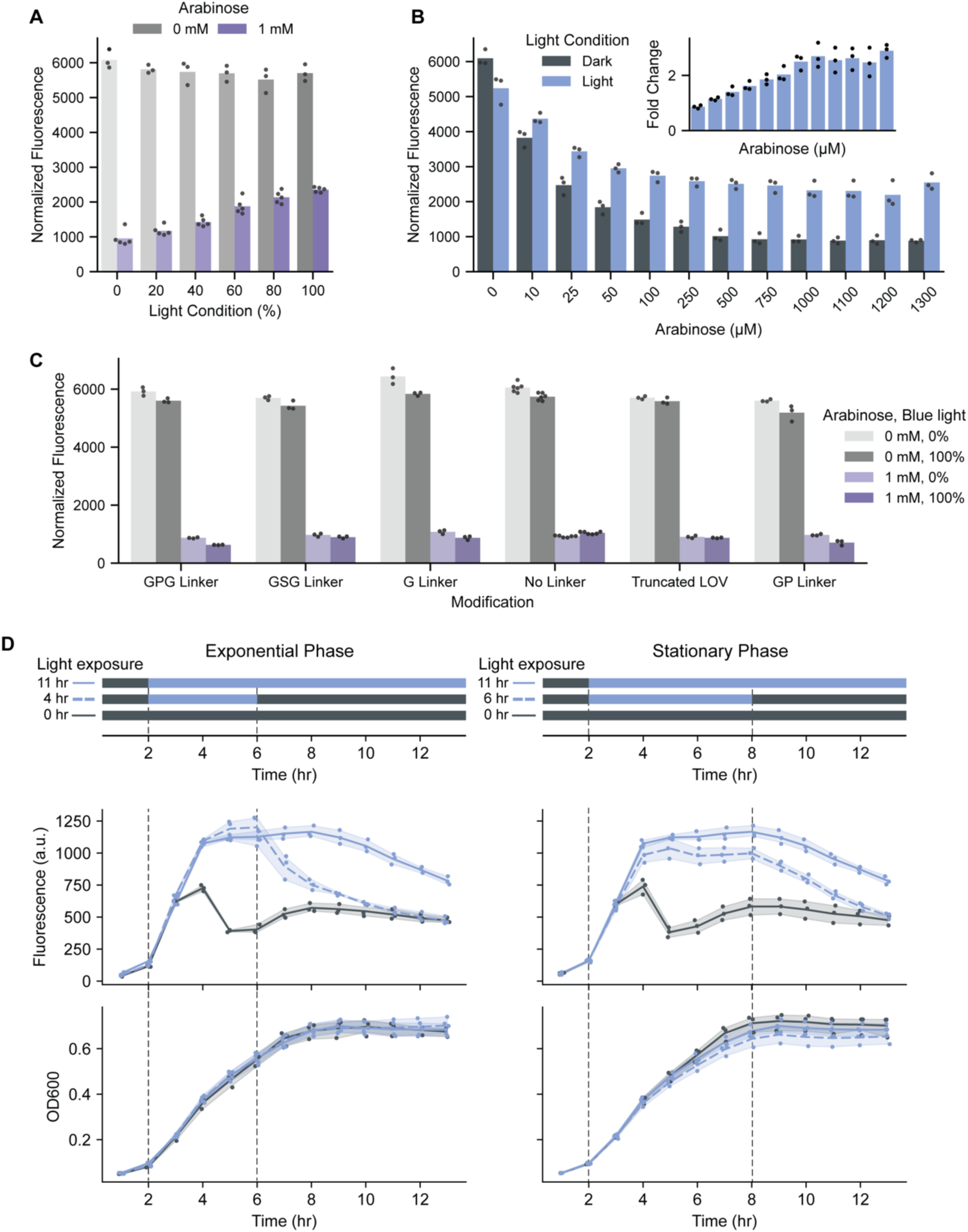
Characterizing, optimizing, and testing dark-deg variant mfLon-LOV-534. (A) Light-dependent proteolytic activity of mfLon-LOV-534. 100% corresponds to 1 mW/cm^2^, with lower intensities scaling linearly. (B) Arabinose-dependent proteolytic activity. Inset shows the dynamic range associated with each arabinose induction condition. (C) Functional assay of mfLon-LOV-534 with different linkers. n ≥ 3 biological replicates for each condition. (D) mfLon-LOV-534 response to 4 or 6 hours of blue light exposure over the course of 13 hours. Targeted protein degradation occurs after blue light removal at the 6- or 8-hour mark, when the culture is in exponential or stationary phase, respectively. mfLon-LOV-534 expression was induced with arabinose (1 mM). Measured fluorescence is of mChartreuse-pdt; expression was induced with IPTG (100 µM). Fluorescence and OD600 were measured every hour. Blue light, 1 mW/cm^2^. n = 3 biological replicates. In all cases, normalized fluorescence corresponds to raw fluorescence divided by optical density at 600 nm.

### Characterization of mfLon-LOV-534 temporal degradation dynamics

To examine the degradation dynamics of mfLon-LOV-534, we conducted time-course experiments during both the exponential and stationary phases of growth. For the exponential phase experiment, mfLon-LOV-534 cultures were first grown in the dark for 2 hours, then exposed to blue light for 4 hours. The light was then turned off, and fluorescence was measured every hour over the subsequent 7 hour period. Upon transfer to dark conditions, fluorescence in the light-exposed culture declined with an estimated half-life of 89 minutes, approaching the baseline level observed in the culture maintained in the dark throughout the experiment (Fig. 5D).

For the stationary phase experiment, mfLon-LOV-534 cultures were grown in the dark for 2 hours and then exposed to blue light for 6 hours, at which point cultures had reached stationary phase (8 hours of growth). Following the transition to dark conditions, we monitored fluorescence for an additional 5 hours. Similar to the exponential phase experiment, the light-exposed culture decreased to fluorescence levels comparable to the dark control (Fig. 5D). Although mChartreuse degradation during stationary phase was slower, with an estimated half-life of approximately 118 minutes, this rate remained faster than the natural decay of the protein observed in stationary phase cells exposed to light for the entirety of the experiment. In cultures with sustained light exposure throughout the experiment, we observed a decrease in fluorescence around 8 hours, likely due to photobleaching of the mChartreuse fluorophore. Despite this unintended effect, targeted degradation of mChartreuse-pdt was achievable in both exponential and stationary phases. Fluorescence levels in both groups converge towards those of the dark-only control. These results confirm that mfLon-LOV-534 exhibits one of the key benefits of light-based enzymatic control, as proteolysis can be achieved independent of growth phase.

## Discussion

In this work, we developed a post-translational optogenetic tool by engineering light-responsive control into the orthogonal *Mesoplasma florum* Lon (mfLon) protease in *E. coli*. Using the SPINE protocol, we generated LOV insertions in mfLon across nearly all codon positions. Our identification of the dark-deg variant mfLon-LOV-534 highlights a fundamental engineering challenge: out of 726 variants tested, only a single site yielded a functional light-responsive protease. This site would likely have been overlooked by rational design alone, as residue 534 lies adjacent to the ATPase domain and does not meet standard insertion criteria. Assuming the complexity exhibited by mfLon extends to other multimeric proteins, these results emphasize the need for high-throughput screening when engineering allosteric control in complex multimeric targets, where functional sites are rare and not readily predictable. Our linker optimization efforts further underscore this point. Substituting the GS linker with combinations differing by as few as one or two amino acids not only failed to improve the dynamic range of mfLon-LOV-534, but it also abolished light responsiveness entirely. This sensitivity raises the possibility that other insertion positions in the library could harbor latent light responsiveness that may be unlocked through systematic linker screening.

We employed a LOV domain variant carrying mutations selected to achieve tighter dark-state caging of the Jα-helix (Fig. S1), and it is important to consider the potential trade-offs of this strategy. While a high dynamic range is a primary goal in optogenetic design, overly tight dark caging could be detrimental to overall tool performance. In the context of mfLon-LOV-534, tighter caging may stabilize the dark conformation and preserve proteolytic activity in the absence of light, a beneficial effect for a dark-active variant. However, it may also promote faster Jα-helix re-docking upon light removal, shortening the effective window of light-induced inhibition and reducing the net impact of blue-light exposure. Furthermore, because mfLon functions as a homohexamer, each protomer carries an independent LOV domain. The six LOV insertions are not obligated to act in concert; stochastic or asynchronous undocking across protomers could dampen the cooperativity of the conformational response and limit the achievable dynamic range. Understanding the degree to which LOV domains within a single hexamer act in tandem will be an important consideration for future optimization.

The expression of mfLon can impose a metabolic burden on the *E. coli* host, which is visible as a reduction in growth (Fig. S8, S11, S13). Optimizing expression levels will be critical for deploying this tool in circuit and metabolic engineering contexts. Further, although we demonstrate that mfLon-LOV-534 is capable of clearing proteins at rates exceeding natural decay in both exponential and stationary phases, its performance remains to be optimized across diverse biological contexts. Future work could evaluate this tool in alternative *E. coli* strains, other microbial chassis, and with target proteins directly relevant to cellular metabolic processes.

The hexameric architecture of mfLon offers an interesting future opportunity to fine-tune proteolytic activity through heterochimeric complex formation. Because mfLon-LOV-534 assembles from six identical protomers each carrying the LOV insertion, the stoichiometric ratio of LOV-bearing to wild-type protomers within a single hexamer could, in principle, be titrated to achieve intermediate levels of light responsiveness. This strategy could also facilitate the discovery of additional functional insertion sites, such as positions that display light-responsive behavior only when a subset of protomers carries the LOV domain rather than all six. More broadly, the orthogonality of the mfLon system is a key advantage since mfLon acts exclusively on the mf-ssrA degron and does not cross-react with endogenous *E. coli* degradation machinery;^31^ it can be deployed as a modular, plug-and-play component without disrupting native proteostasis. The orthogonality and tunability of mfLon-LOV-534 make it an interesting tool for post-translational control in synthetic biology, and this work establishes a generalizable approach for engineering allosteric regulation in complex enzymes.

## Methods

### Strains and plasmids

All plasmids were constructed using *E. coli* DH5α, while functional assays for the mfLon-LOV library and subsequent characterization experiments were performed in the *E. coli* K-12 strain BW25113.^40^ To generate mfLon-LOV variants we utilized pZA16mflon (Addgene #75439) and pBbS5c-mCherry-AsLOV2*(543) (Addgene #202712). AsLOV2*(543) has a C-terminal 3 amino acid truncation and mutations L493V, H519R, V520L, D522G, G528A, E537F, N538Q, and D540A for tighter dark caging (Fig. S1). When constructing individual variants, the sequence for AsLOV2*(543), which we refer to as ‘LOV’ in this manuscript for simplicity, was amplified and inserted immediately following the chosen insertion sites via Golden Gate assembly.^41^

We developed the base IPTG-inducible expression plasmid, pBbS5c-mChartreuse, by replacing the mRFP1 sequence of the BglBrick backbone pBbS5c-mRFP1^42^ with mChartreuse. The fluorophore mChartreuse is a modified green fluorescent protein with enhanced photostability and brightness relative to sfGFP.^43^ The sequence for mChartreuse was amplified from pNF02-mChartreuse (Addgene #219397) and used to replace the mCherry sequence in pBbS5c-mCherry-pdt3 via Golden Gate assembly. For mfLon functional assays, we constructed a reporter that features mChartreuse fused to the modified degradation tag, pdt3 (amino acid sequence: AANKNEENTNEVPTFMLNAGQANRRRV) from Cameron et al.^31^ The pdt3 sequence was sourced from plasmid pZE27MC3 (Addgene #75464). We constructed pBbS5c-mChartreuse-pdt3 by inserting a synthetic double-stranded DNA fragment (gBlock, Integrated DNA Technologies) encoding the pdt3 degron into the pBbS5c-mChartreuse backbone via Golden Gate assembly. To screen the mfLon-LOV library, the lacUV5 promoter of this reporter was replaced with the constitutively active W7 promoter (TTATCAAAAAGAGTATTGAAATAAAGTCTAACCTATAG GAAGATTACAGCCATCGAG AGGGACACGGCGAA), generating pW7-mChartreuse-pdt3. All other experiments use the IPTG inducible version of the mChartreuse-pdt reporter. For all functional characterization, the respective reporter plasmids were co-transformed with the corresponding mfLon-LOV variant plasmids into *E. coli* BW25113.

### Comprehensive insertion of LOV into mfLon to generate mfLon-LOV library

To design and construct a comprehensive insertional library of LOV in mfLon, we used the saturated programmable insertion engineering (SPINE) technique.^26^ The purpose of SPINE is to generate an unbiased library that covers every possible in-frame insertion of a domain into a gene of interest. For the mfLon-LOV library, the gene of interest was *mfLon*, contained in pZA16mflon, and the insertion domain was AsLOV2*(543) (LOV), amplified from pBbS5c-mCherry-AsLOV2*(543), as described above. Using the SPINE computational pipeline, we split *mfLon* into 13 fragments, each comprised of ∼60 amino acids. These fragments were then used to generate synthesizable oligo sequences flanked by fragment-specific barcodes and external BsmBI cut sites. Oligo pools were synthesized by Twist Bioscience. Within each fragment, each oligo contains a genetic handle with internal BsaI cut sites placed after each codon; collectively, the oligo library spans all possible insertion positions. Oligo sub-libraries for each of the 13 fragments were amplified using their respective barcodes. For every fragment, the SPINE approach was also used to generate a pair of primers to amplify a partial sequence of *mfLon* that excludes the corresponding fragment region and adds external BsmBI cut sites that are complementary to those on the corresponding fragment oligos. Using the Golden Gate method, these components produced a comprehensive mfLon library in which a BsaI cut site occupies one codon position per construct, thereby covering all possible in-frame positions. We refer to the resulting library as our first intermediate library.

Primers were designed to amplify a chloramphenicol cassette with the resulting amplicon containing external BsaI cut sites flanked by internal BsmBI cut sites. The chloramphenicol acetyl transferase (*cat*) gene was sourced and amplified from pBbS5c-mRFP1.^44^ Using the Golden Gate method, the cassette replaced the genetic handle in our first intermediate library and served as a positive selection marker for successful oligo integration into *mfLon*. We refer to the resulting library after cassette insertion as our second intermediate library.

Next, the insertional domain, LOV, was amplified and introduced into the library using the Golden Gate method by excising the internal BsmBI cut sites of the chloramphenicol cassette in the second intermediate library. This resulted in the final insertion library of mfLon-LOV variants.

At all steps in the library generation process, colony-forming unit (CFU) counts were performed on plates containing appropriate antibiotics to confirm that the number of transformants exceeded the expected number of variants by one order of magnitude. For example, after inserting oligo subpools of the first fragment, the resulting library should contain 60 variants; thus, we verified that CFU counts were >600. Resulting amplicons throughout the process were gel purified using the Zymo Gel DNA Extraction Kit. All PCR amplifications were conducted using KAPA HiFi HotStart ReadyMix 2x (Roche).

### Bacterial cell culture and blue-light stimulation

Unless stated otherwise, bacteria were cultured in Luria Broth (LB) supplemented with appropriate antibiotics for plasmid maintenance and grown at 37 °C with shaking at 200 rpm. The reporter plasmids pBbS5c-mCherry-pdt3 and pW7-mChartreuse-pdt3 contain chloramphenicol resistance genes and the SC101 origin of replication. Plasmids containing wild-type mfLon or mfLon-LOV variants contain a carbenicillin resistance gene and origin of replication p15A. Antibiotic concentrations used for plasmid maintenance were 100 µg/mL for carbenicillin and 25 µg/mL for chloramphenicol. Light-exposure experiments on rationally designed mfLon-LOV constructs were conducted in an optoWELL-96 device (Opto Biolabs) using 460 nm blue light. Light-exposure experiments on the mfLon-LOV library were conducted on an optoWELL-24 device (Opto Biolabs) using 460 nm blue light. Unless stated otherwise, overnight cultures of the reference strain and mfLon-LOV library were diluted 1:100, plated, and pre-cultured in the dark for 2 hours before light exposure. IPTG (100 µM) and arabinose induction were added during the initial dilution step for cultures receiving these inducers. Unless otherwise stated, light exposure lasted 4 hours at a light intensity of 1 mW/cm^2^. Green fluorescence (excitation 480 nm, emission 515 nm), and optical density (OD at 600 nm) readings were taken using a BioTek Synergy H1m plate reader (BioTek, Winooski, VT) at the end of incubation.

For sorting, cells were spun down and diluted 1:1000 in 1x phosphate-buffered saline (PBS). Cells collected during sorting were resuspended in LB, antibiotics were added for plasmid maintenance, and the cells were grown overnight at 30 °C to avoid overgrowth, per the Coyote-Maestas et al. methods.^26^

### Cell sorting

Fluorescence-activated cell sorting (FACS) was performed on a Sony SH800S using a 100 µm sorting chip. Singlet events were gated using side-scatter width (SSC-W) versus side-scatter area (SSC-A) to exclude doublets or aggregates. Fluorescence gates were defined using a reference strain co-transformed with pW7-mChartreuse-pdt3 and pZA16mflon, in which mfLon expression was controlled by an arabinose-inducible promoter. The low fluorescence gate (-mChartreuse) was determined using the arabinose-induced (1 mM) reference strain, which degrades mChartreuse-pdt3. The high fluorescence gate (+mChartreuse) was determined using the uninduced reference strain. The population of cells co-transformed with the mfLon-LOV library and reporter plasmid pW7-mChartreuse-pdt3 is referred to as the naïve library. The naïve library and sorted populations grown under either dark or light conditions were analyzed using the same settings defined above. For each condition and gate, cells were sorted in ultrapurity mode to collect 200,000 events.

We conducted a sorting enrichment campaign where we defined four bins, corresponding to both the light condition the culture was grown in (Dark/Light) and the gate used to sort the population (-/+mChartreuse). In the FACS enrichment campaign, the naïve library was initially grown in the dark or under blue light and sorted into −/+mChartreuse, resulting in four bins: (i) Dark/-mChartreuse, (ii) Dark/+mChartreuse, (iii) Light/-mChartreuse, and (iv) Light/+mChartreuse. After sorting, the cultures from each bin were grown overnight in LB medium containing carbenicillin and chloramphenicol for plasmid maintenance, and the overnight cultures were stored as glycerol stocks; these cultures were designated as enrichment round 1. For subsequent enrichment rounds, overnights from the previous sorting round, grown directly from the sorting round or from the glycerol stock, were diluted and plated under the same light conditions, and sorted using the same gate. This process was repeated for a total of six rounds.

### Sequencing and alignment

Nanopore sequencing was performed on mfLon-LOV library plasmids isolated by miniprep following FACS sorting (or before any sorting, for the naïve library kept in the dark or exposed to blue light). Prior to sequencing (Plasmidsaurus), mfLon-LOV library samples were prepared using the ZymoPURE II Plasmid Midiprep Kit (Zymo Research, Cat# D4200) and digested with LguI to linearize the plasmid. DNA concentration was measured using the broad range Qubit dsDNA Quantification Assay Kit (Thermo Fisher Scientific, Cat# Q32850), and samples were diluted to Plasmidsaurus standards. Since the library generation process is imperfect, plasmids with no oligo insertions can still be present. Digestion with LguI enabled linearization of the DNA without amplifying the library, avoiding PCR-based library amplification and potential distortion of variant representation. We used the Plasmidsaurus “huge” amplicon sequencing option (up to 12,000 raw reads) for library intermediates containing internal BsaI cut sites or chloramphenicol cassettes (samples diluted to 40 µL at 50 ng/µL). To sequence sorted libraries after each of the six rounds of enrichment, as well as the naïve library, we placed a custom sequencing order for 2 Gb reads (samples diluted to 40 µL at 55.7 ng/µL).

Sequencing data were analyzed using a custom Python script (https://gitlab.com/dunloplab/spine-sequencing-analysis) we developed that combined multiple packages to detect LOV insertion locations. First, NanoFilt^45^ was used to filter nanopore sequencing reads by a minimum Phred score; our threshold was set to 10. To track reads lost due to filtering, we calculated the number of reads before and after filtering. Filtered reads were aligned to the wild-type mfLon sequence using either minimap2^46^ or NGMLR.^47^ Alignments were processed using SAMtools^48^ to generate .bai index files. Reads were split by alignment orientation (forward/reverse) and converted back to FASTQ format. Reverse-oriented sequences were transformed into their reverse complement using seqtk (https://github.com/lh3/seqtk.git). To find raw counts of LOV insertions at specific codons in the *mfLon* gene, we provided 2FAST2Q^49^ the resulting FASTQ files containing aligned forward and reverse-corrected reads. Briefly, 2FAST2Q works by parsing every read in the FASTQ files, attempting to match a search sequence at any position within each read. In our case, the search sequence was the first 12 base pairs of the LOV insertion, including the GS linker introduced through the SPINE process (AGCGGGTTGGCT). Once 2FAST2Q found a match for the search sequence, the 18 bases upstream of the search sequence were extracted and matched against a provided feature library. The feature library matched the 18-base-pair upstream sequence to the codon position. 2FAST2Q counted the number of instances of each feature in the FASTQ files. Overall, the analysis identified the location of LOV insertions for all sequence reads.

### Enrichment analysis of sequencing data

First, we performed data normalization and fold-change calculations. We conducted an enrichment analysis to visualize shifts in insertion site enrichment or depletion and to map functional categories based on these shifts. The resulting data from the sequencing and alignment pipeline is an array (*C*) of length *N*, where *N* is the number of codons in mfLon, describing the raw counts of LOV insertion at each codon position. Raw sequencing counts of the naïve library and the binned populations were normalized by their sequencing depth. The resulting array, NormCount, describes the frequency of a variant for a particular bin.

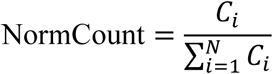

Enrichment for all four bins was calculated as the *Log*_2_fold-change relative to the naïve library. To avoid division by 0, a pseudocount (*pc* = 1) was applied.

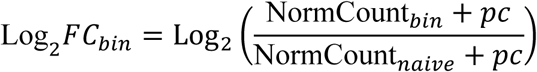

Thus, if an insertion site is overrepresented in a bin relative to the naïve library, the corresponding value is positive; if depleted, it is negative.

Next, we made quadrant assignments for variant category prediction. From the fold-changes, we defined a differential metric to compare relative enrichment between –mChartreuse and +mChartreuse within the same light conditions (Dark/Light):

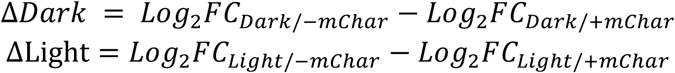

To assign potential functional significance to the variants, we mapped each insertion into a Cartesian coordinate system using its ΔDark and ΔLight values.

### Construction of linker variants

Variants of mfLon-LOV-534 with altered linkers were constructed directly from mfLon-LOV-534 through Golden Gate assembly. Primers were designed to amplify the backbone and the LOV insert while selectively preserving residues from the glycine residue in the original GS linker and include new residues in the primer overhangs. The amplified products were then assembled using the Golden Gate method with the respective type IIS restriction enzyme (BsaI or BsmBI). The linker sequences are as follows: 5’-GGGCCTGGG-3’ (GPG), 5’-GGGAGCGGG-3’ (GSG), 5’-GGG-3’ (G), 5’-CCAGGG-3’ (GP). All linkers flanked the N- and C-terminal ends of the LOV domain. The GP linker is not palindromic, and it was arranged as follows: PG-LOV-GP.

To create mfLon-LOV-534 with no linkers (mfLon-LOV-534-NL), the backbone of the mfLon-LOV-534 plasmid was amplified, excluding the LOV domain flanked by GS linkers. We amplified a LOV insert from the mfLon-LOV-534 plasmid, excluding the GS linkers flanking the sequence. The LOV domain was inserted into the backbone via Golden Gate assembly.

To create the N-terminal LOV truncated mfLon-LOV-534 variant, we amplified the mfLon-LOV-534-NL to exclude the first 3 N-terminal amino acids of the LOV domain (LAT). We designed the primers to have a BsmBI overhang, which we self-ligated post-amplification.

### Protein visualization and rational design metrics

Protein structure visualizations and solvent-accessible surface area calculations were performed using ChimeraX.^50–52^ Solvent-accessible surface area threshold was set to >30Å^%^ based on methods from Dagliyan et al.^21^ The evolutionary conservation of residues was estimated using ConSurf^53^ and visualized with ChimeraX. ProDomino predictions were executed following the package instructions located in GitHub (https://github.com/Niopek-Lab/ProDomino).^27^

## Acknowledgements

We thank Dr. Winston Timp for initial guidance on sequencing analysis and Eric South for his help using FACS. This work was supported by National Science Foundation grant CBET-2422796 and National Institutes of Health grant R01AI102922. Lily Fenton received support from the NSF NRT on Biological Feedback Control, DGE-2244366.

## References

1. Lindner, F. & Diepold, A. Optogenetics in bacteria – applications and opportunities. FEMS Microbiol Rev 46, fuab055 (2022).

2. Huang, H. & Dunlop, M. J. Single-cell analysis and control of microbial systems using optogenetics. Current Opinion in Microbiology 89, 102702 (2026).

3. Wegner, S. A., Barocio-Galindo, R. M. & Avalos, J. L. The bright frontiers of microbial metabolic optogenetics. Current Opinion in Chemical Biology 71, 102207 (2022).

4. Wei, J. & Jin, F. Illuminating bacterial behaviors with optogenetics. Current Opinion in Solid State and Materials Science 26, 101023 (2022).

5. Hoffman, S. M., Tang, A. Y. & Avalos, J. L. Optogenetics Illuminates Applications in Microbial Engineering. Annual Review of Chemical and Biomolecular Engineering 13, 373–403 (2022).

6. Ochoa-Fernandez, R. et al. Optogenetic control of gene expression in plants in the presence of ambient white light. Nat Methods 17, 717–725 (2020).

7. Castillo-Hair, S. M., Baerman, E. A., Fujita, M., Igoshin, O. A. & Tabor, J. J. Optogenetic control of Bacillus subtilis gene expression. Nat Commun 10, 3099 (2019).

8. Zhao, E. M. et al. Optogenetic Amplification Circuits for Light-Induced Metabolic Control. ACS Synth. Biol. 10, 1143–1154 (2021).

9. Gil, A. A. et al. Optogenetic control of protein binding using light-switchable nanobodies. Nat Commun 11, 4044 (2020).

10. Lugagne, J.-B., Blassick, C. M. & Dunlop, M. J. Deep model predictive control of gene expression in thousands of single cells. Nat Commun 15, 2148 (2024).

11. Blassick, C. M., Moghimianavval, H., Jafarbeglou, F., Lugagne, J.-B. & Dunlop, M. J. Dynamic heterogeneity in an E. coli stress response regulon mediates gene activation and antimicrobial peptide tolerance. 2024.11.27.625634 Preprint at 10.1101/2024.11.27.625634 (2025).

12. Tabor, J. J., Levskaya, A. & Voigt, C. A. Multichromatic Control of Gene Expression in Escherichia coli. Journal of Molecular Biology 405, 315–324 (2011).

13. Sheets, M. B., Wong, W. W. & Dunlop, M. J. Light-Inducible Recombinases for Bacterial Optogenetics. ACS Synthetic Biology 9, 227–235 (2020).

14. Nath, K. & Koch, A. L. Protein Degradation in *Escherichia coli*. Journal of Biological Chemistry 246, 6956–6967 (1971).

15. Tague, N., Coriano-Ortiz, C., Sheets, M. B. & Dunlop, M. J. Light inducible protein degradation in E. coli with the LOVdeg tag. eLife 12, (2024).

16. Komera, I. et al. Bifunctional optogenetic switch for improving shikimic acid production in E. coli. Biotechnology for Biofuels and Bioproducts 15, 1–14 (2022).

17. Chen, Y. et al. Design principles for optogenetic-based targeted protein degradation. Synth Syst Biotechnol 12, 255–264 (2025).

18. Li, Z. et al. De novo designed protein guiding targeted protein degradation. Nat Commun 16, 6598 (2025).

19. Gur, E. & Sauer, R. T. Evolution of the ssrA degradation tag in Mycoplasma: Specificity switch to a different protease. Proceedings of the National Academy of Sciences 105, 16113–16118 (2008).

20. Mathony, J., Aschenbrenner, S., Becker, P. & Niopek, D. Dissecting the Determinants of Domain Insertion Tolerance and Allostery in Proteins. Advanced Science 10, 2303496 (2023).

21. Dagliyan, O., Dokholyan, N. V. & Hahn, K. M. Engineering proteins for allosteric control by light or ligands. Nat Protoc 14, 1863–1883 (2019).

22. Lee, J. et al. Surface Sites for Engineering Allosteric Control in Proteins. Science 322, 438–442 (2008).

23. Dagliyan, O. et al. Engineering extrinsic disorder to control protein activity in living cells. Science 354, 1441–1444 (2016).

24. Reynolds, K. A., McLaughlin, R. N. & Ranganathan, R. Hot Spots for Allosteric Regulation on Protein Surfaces. Cell 147, 1564–1575 (2011).

25. Zhu, L., McNamara, H. M. & Toettcher, J. E. Light-switchable transcription factors obtained by direct screening in mammalian cells. Nat Commun 14, 3185 (2023).

26. Coyote-Maestas, W., Nedrud, D., Okorafor, S., He, Y. & Schmidt, D. Targeted insertional mutagenesis libraries for deep domain insertion profiling. Nucleic Acids Res 48, e11 (2020).

27. Wolf, B. et al. Rational engineering of allosteric protein switches by in silico prediction of domain insertion sites. 2024.12.04.626757 Preprint at 10.1101/2024.12.04.626757 (2024).

28. Carrasco-López, C. et al. Development of light-responsive protein binding in the monobody non-immunoglobulin scaffold. Nat Commun 11, 4045 (2020).

29. Guntas, G. et al. Engineering an improved light-induced dimer (iLID) for controlling the localization and activity of signaling proteins. Proceedings of the National Academy of Sciences of the United States of America 112, 112–117 (2015).

30. Fraikin, N., Couturier, A., Mercier, R. & Lesterlin, C. A palette of bright and photostable monomeric fluorescent proteins for bacterial time-lapse imaging. Science Advances 11, eads6201 (2025).

31. Cameron, D. E. & Collins, J. J. Tunable protein degradation in bacteria. Nat Biotechnol 32, 1276–1281 (2014).

32. Abramson, J. et al. Accurate structure prediction of biomolecular interactions with AlphaFold 3. Nature 630, 493–500 (2024).

33. Wlodawer, A., Sekula, B., Gustchina, A. & Rotanova, T. V. Structure and the mode of activity of Lon proteases from diverse organisms. J Mol Biol 434, 167504 (2022).

34. Coscia, F. & Löwe, J. Cryo-EM structure of the full-length Lon protease from Thermus thermophilus. FEBS Letters 595, 2691–2700 (2021).

35. Paysan-Lafosse, T. et al. The Pfam protein families database: embracing AI/ML. Nucleic Acids Res 53, D523–D534 (2025).

36. Abramson, J. et al. Accurate structure prediction of biomolecular interactions with AlphaFold 3. Nature 630, 493–500 (2024).

37. Liu, M. et al. OptoLacI: optogenetically engineered lactose operon repressor LacI responsive to light instead of IPTG. Nucleic Acids Research 52, 8003–8016 (2024).

38. Nadler, D. C., Morgan, S.-A., Flamholz, A., Kortright, K. E. & Savage, D. F. Rapid construction of metabolite biosensors using domain-insertion profiling. Nat Commun 7, 12266 (2016).

39. Mathony, J. & Niopek, D. Enlightening Allostery: Designing Switchable Proteins by Photoreceptor Fusion. Advanced Biology 5, 2000181 (2021).

40. Baba, T. et al. Construction of Escherichia coli K-12 in-frame, single-gene knockout mutants: the Keio collection. Mol Syst Biol 2, MSB4100050 (2006).

41. Engler, C., Gruetzner, R., Kandzia, R. & Marillonnet, S. Golden Gate Shuffling: A One-Pot DNA Shuffling Method Based on Type IIs Restriction Enzymes. PLOS ONE 4, e5553 (2009).

42. Lee, T. S. et al. BglBrick vectors and datasheets: A synthetic biology platform for gene expression. J Biol Eng 5, 12 (2011).

43. Pédelacq, J.-D., Cabantous, S., Tran, T., Terwilliger, T. C. & Waldo, G. S. Engineering and characterization of a superfolder green fluorescent protein. Nat Biotechnol 24, 79–88 (2006).

44. Lee, T. S. et al. BglBrick vectors and datasheets: A synthetic biology platform for gene expression. Journal of Biological Engineering 5, 12 (2011).

45. De Coster, W., D’Hert, S., Schultz, D. T., Cruts, M. & Van Broeckhoven, C. NanoPack: visualizing and processing long-read sequencing data. Bioinformatics 34, 2666–2669 (2018).

46. Li, H. New strategies to improve minimap2 alignment accuracy. Bioinformatics 37, 4572–4574 (2021).

47. Sedlazeck, F. J. et al. Accurate detection of complex structural variations using single-molecule sequencing. Nat Methods 15, 461–468 (2018).

48. Danecek, P. et al. Twelve years of SAMtools and BCFtools. Gigascience 10, giab008 (2021).

49. Bravo, A. M., Typas, A. & Veening, J.-W. 2FAST2Q: a general-purpose sequence search and counting program for FASTQ files. PeerJ 10, e14041 (2022).

50. Goddard, T. D. et al. UCSF ChimeraX: Meeting modern challenges in visualization and analysis. Protein Science 27, 14–25 (2018).

51. Pettersen, E. F. et al. UCSF ChimeraX: Structure visualization for researchers, educators, and developers. Protein Science 30, 70–82 (2021).

52. Meng, E. C. et al. UCSF ChimeraX: Tools for structure building and analysis. Protein Science 32, e4792 (2023).

53. Ashkenazy, H. et al. ConSurf 2016: an improved methodology to estimate and visualize evolutionary conservation in macromolecules. Nucleic Acids Res 44, W344–W350 (2016).

