## Supplementary Information for "Engineering a light-responsive, orthogonal Lon protease in *E. coli* for targeted protein degradation"

#### Supplementary Tables

**Table S1.** Insertion site list, rationale for inclusion, and ΔDark-ΔLight prediction.

| Insertion site | Rational for inclusion | ΔDark-ΔLight Prediction |
|---|---|---|
| **80** | ΔDark-ΔLight, ProDomino | Light-deg |
| **81** | Rational | Inert |
| **82** | Rational | Inert |
| **83** | Rational | Inert |
| **104** | Rational | Inert |
| **105** | Rational | Inert |
| **118** | ProDomino | Inert |
| **119** | ΔDark-ΔLight | Inert |
| **122** | ProDomino | Inert |
| **132** | ΔDark-ΔLight | Light-deg |
| **144** | ProDomino | Inert |
| **190** | ΔDark-ΔLight | Light-deg |
| **191** | ΔDark-ΔLight | Nonfunctional |
| **202** | ΔDark-ΔLight | Dark-deg |
| **205** | ΔDark-ΔLight | Inert |
| **218** | ΔDark-ΔLight | Light-deg |
| **301** | ΔDark-ΔLight | Light-deg |
| **306** | ΔDark-ΔLight | Nonfunctional |
| **471** | ΔDark-ΔLight | Light-deg |
| **520** | ΔDark-ΔLight | Dark-deg |
| **534** | ΔDark-ΔLight | Dark-deg |
| **537** | ΔDark-ΔLight | Nonfunctional |
| **543** | ΔDark-ΔLight | Dark-deg |
| **565** | ΔDark-ΔLight | Dark-deg |
| **572** | ΔDark-ΔLight | Dark-deg |
| **671** | ΔDark-ΔLight | Nonfunctional |
| **710** | ΔDark-ΔLight | Light-deg |
| **711** | Rational | On axis |
| **712** | Rational | Nonfunctional |
| **713** | Rational | Nonfunctional |

### Supplementary Figures

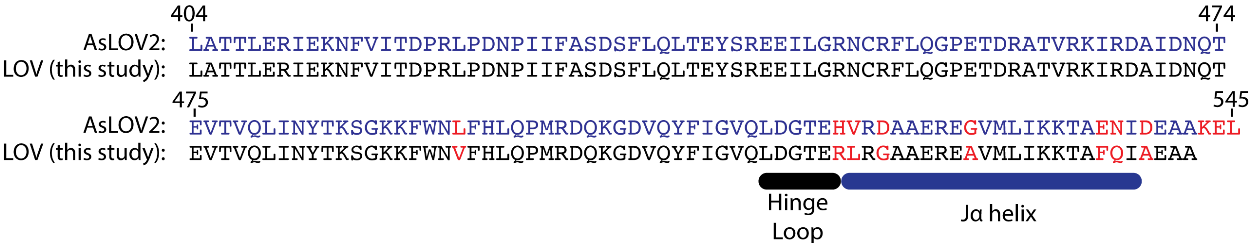


**Figure S1.** Comparison between wild-type AsLOV2 amino acid sequence and mutations of LOV used in this study for tighter dark caging.

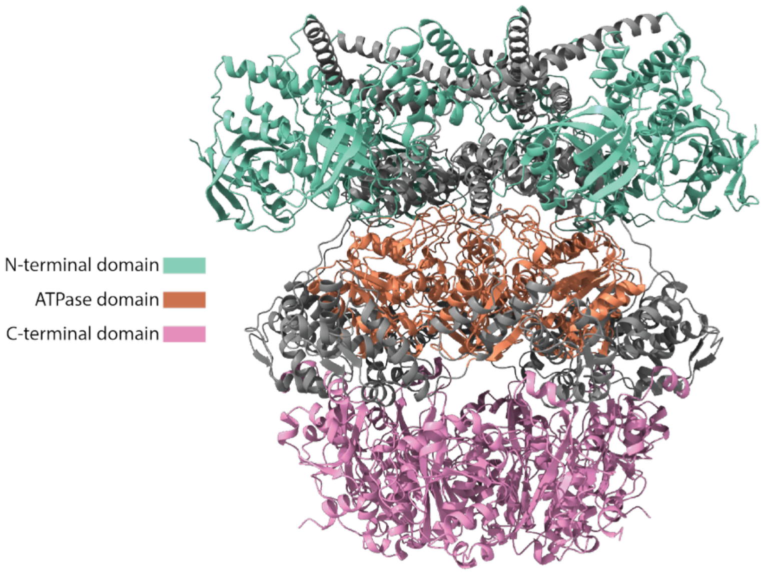


**Figure S2.** Highlighted are the functional regions of the mfLon protein. The N-terminal domain ranges from residues 1-211, the ATPase domain ranges from residues 366-503, and the C-terminal domain ranges from residues 584-787. Regions were identified using the Pfam protein database that utilizes homology to highlight functional regions.

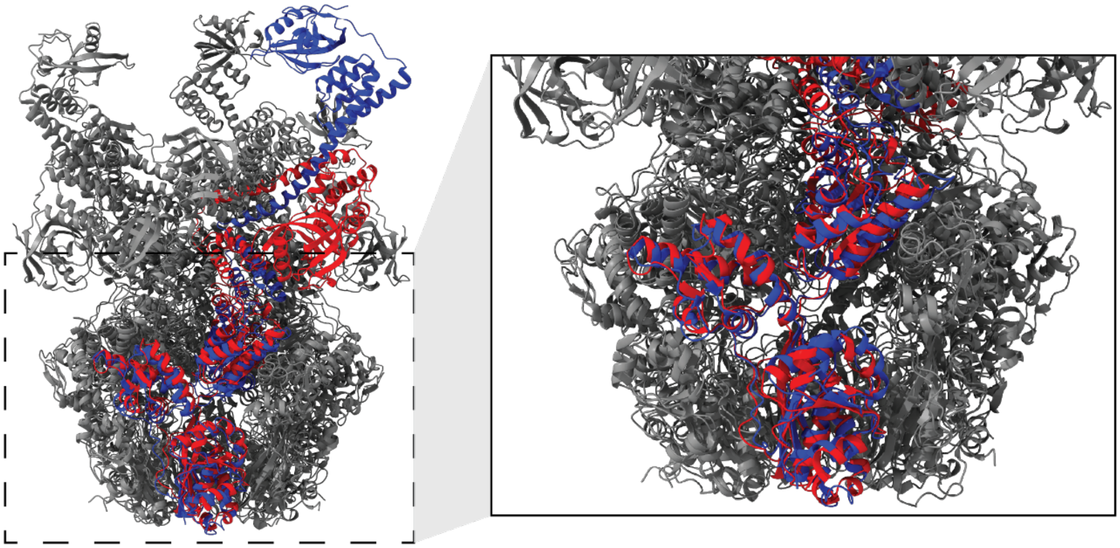

**Figure S3.** Alignment of *Thermus thermophilus* Lon (ttLon, PDB: 7P6U) with the AlphaFold3-predicted mfLon structure reveals similarities. Blue corresponds to ttLon; red corresponds to mfLon. Alignment performed using the ChimeraX matchmaker function. On the left, the full structures are aligned. On the right, the ATPase and C-terminal domains are highlighted. The N-terminal domains are not perfectly aligned due to homolog variability.

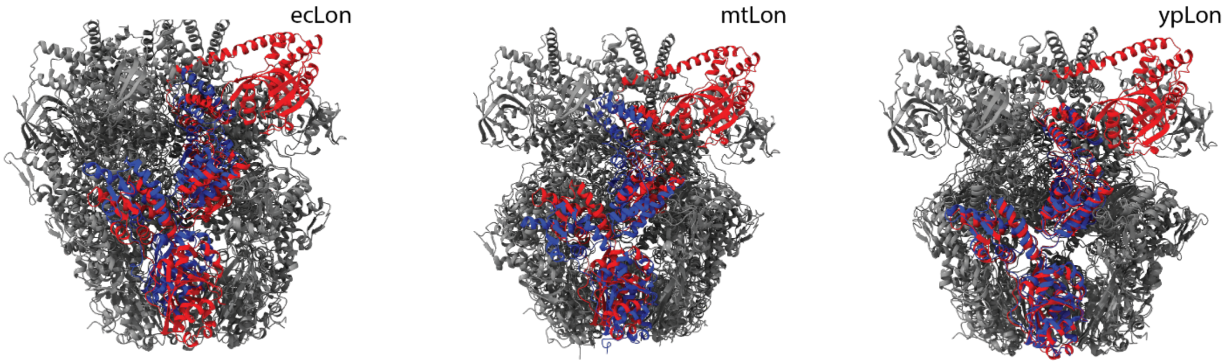


**Figure S4.** Predicted mfLon structure aligned with *Escherichia coli* ecLon (PDB: 6U5Z), *Meiothermus taiwanensis* mtLon (PDB: 4YPL), and *Yersinia pestis* ypLon (PDB: 6ON2). Blue corresponds to the respective homolog; red corresponds to mfLon. Alignment performed using ChimeraX matchmaker function.

```
1            11           21           31           41
MSKKIKLPIF   QIRGSFIVPG   IKENLEVGRK   NTLASVNYAI   KNSNNQMIAI
eeeeeebebb   ebeebbbbeb   bbbebebeee   ebeebbeebe   eeeeeebbbb
       f             f

51           61           71           81           91
PQIDASVEKP   EFSDLHEFGI   LIDFEVIKEW   KDNSLTISTN   PIQRCKVISF
beeeeeeeee   eeeebeebbb   bbebeebbee   eeeebebbbe   beeebebeeb
 f                   s                                   f

101          111          121          131          141
FENEDQVPYA   EVELIESIND   FSDEELKELI   EKISDAIKTK   ASLVTKQIKQ
eeeeeeebeb   ebeebeeeee   eeeeebeebb   eebeeebeeb   beeeeeebee

151          161          171          181          191
LISGESDDLS   LAFDSIMFKL   APSKILTNPE   YITSPSLKTR   WSIIEKIIFA
ebeeeeeeee   ebbebbbeeb   eebeeeeeee   bbeeeeeeee   beebbeebee
                f                    f    s       f

201          211          221          231          241
EDGIITRNAE   SIDAARQKNE   IEQELNHKLK   EKMDKQQKEY   YLREKMRIIK
eeebbbeebe   bbebbbebbe   beeebeeebe   eebeeeeeee   bbeeebeebe
f                                            ff        sfff   s

251          261          271          281          291
DELEDEDDSD   DSSLEKYKER   LAKEPFPEEV   KRKIMASIKR   VEALQSGTPE
eeeeeeeeee   eeebeebeee   beeeebeeeb   eeebeeebee   beeeeeeeee
 ff                                          f  f            f

301          311          321          331          341
WNTEKNYIDW   MMSIPWWEET   EDLTDLKYAK   KILDKHHYGM   KKVKERIIEY
bebbeebbeb   bbebeeeeee   eeebebeebe   ebbeeeeeeb   eebeeebbeb
    f s          ff           f   s      s  ff  s      f fs f

351          361          371          381          391
LAVKTKTKSL   KAPIITLVGP   PGVGKTSLAK   SIAEAVGKNF   VKVSLGGVKD
bbbeeeeeee   eeebbbbbbe   eeeeeebebe   bbbebbeeeb   bebbbebbee
 ss  f          s    sf   ffffffsf     s s    f      f ssfs  f

401          411          421          431          441
ESEIRGHRKT   YVGSMPGRII   QTMKRAKVKN   PLFLLDEIDK   MASDHRGDPA
ebebeeeeeb   bbbbbebebb   ebbeeeeeee   ebbbbeebee   beeebeeeeb
fsfsfffffs   s s  fsf              f   f    ff ff      f  fff

451          461          471          481          491
SAMLEVLDPE   QNKEFSDHYI   EEPYDLSQVM   FIATANYPED   IPEALYDRME
ebbbebbeee   eeeebeeebb   ebebebbebb   bbbbbeeeee   beeebeeebe
fs sfssfff   ff  s ff         fss      s  ssf       sf  s ffsf

501          511          521          531          541
IINLSSYTEI   EKVKIAQDYL   VPKAIEQHEL   TSEEISFTEG   AINEIIKYYT
bbebeeeeee   eebebbeeeb   beeeeeeeee   eeeebebeee   bbeebbeebb
 s   fff     ff  ss   s      f    f                         ss

551          561          571          581          591
REAGVRQLER   HINSIIRKYI   VKNLNGEMDK   IVIDEKQVND   LLGKRIFDHT
eeebbeebee   ebeebbeebb   eeebeeeeee   eebeeeebee   beeeeeeeee
fffssf sff         ff                                ff
```

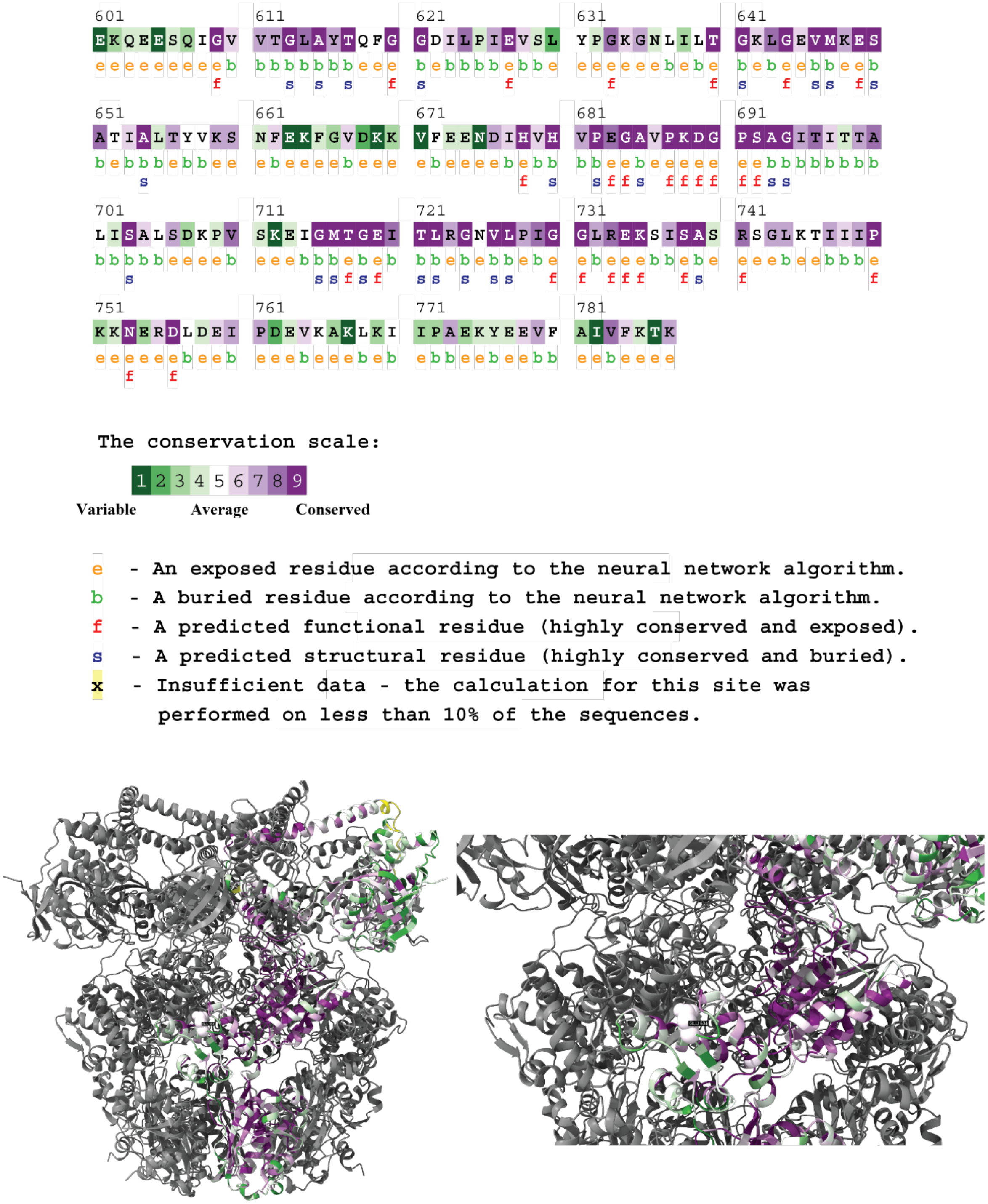


**Figure S5.** Evolutionary conservation of mfLon residues estimated using ConSurf (DOI: 10.1093/nar/gkw408).

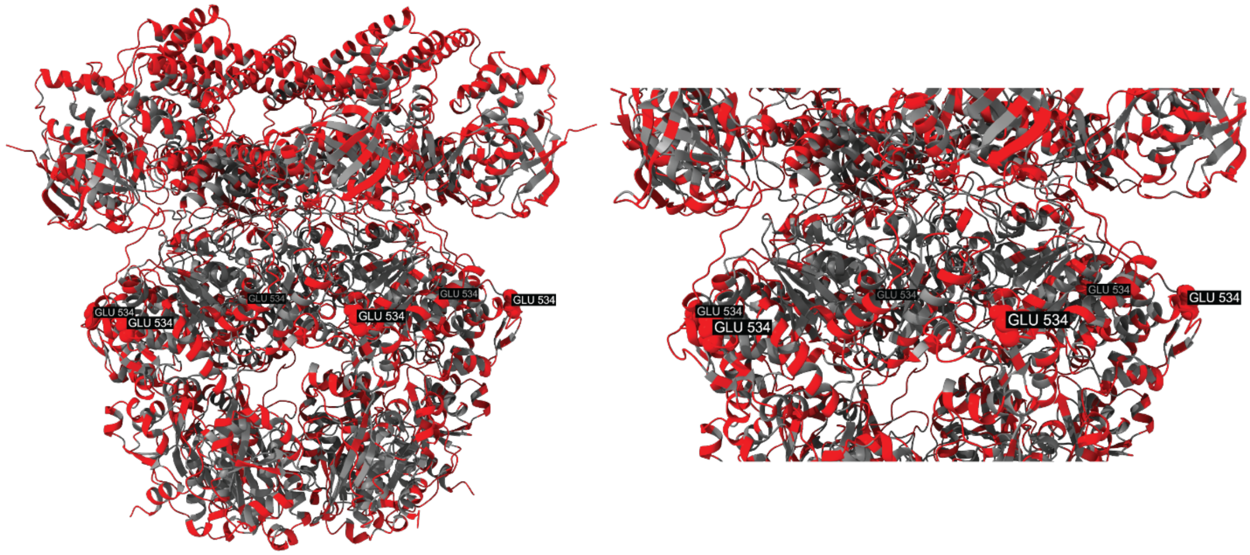


**Figure S6.** The solvent-accessible surface area of the mfLon structure was calculated in ChimeraX. Highlighted in red are residues that had values >30Å$^2$, a cutoff based on the methods of Dagliyan et al. (DOI: 10.1038/s41596-019-0165-3) for rationally finding functional insertion sites.

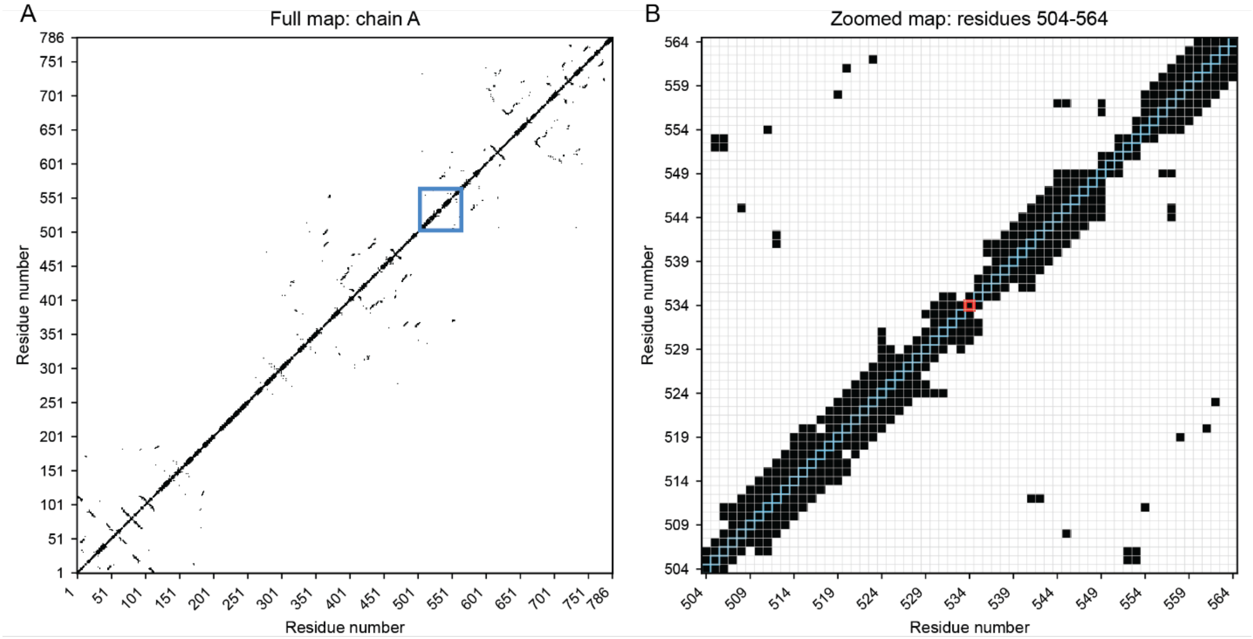


**Figure S7.** Using the mfLon PDB file we created a residue contact map in which residues <7Å from each other are labeled in black. The diagonal represents a residue's interaction with itself, which will always meet the threshold. (A) Contact map of the full mfLon sequence. Regions of clustered black grids that run perpendicular to the diagonal line are locally interacting residues. Regions of clustered black grids are otherwise nonlocal interacting residues. The highlighted region (blue) encompasses residues 504 through 564. (B) Zoomed in contact map from residues 504 through 564. Highlighted in blue is the diagonal. Highlighted in red is residue 534. Residue 534 has no local and few nonlocal interactions.

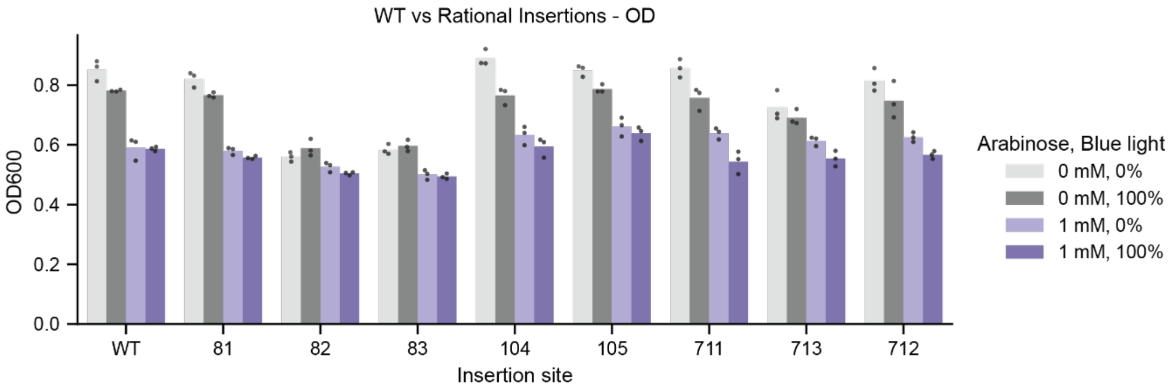


**Figure S8.** Optical density at 600 nm of the rationally selected insertion sites. WT corresponds to strains co-transformed with wild-type mfLon and mChartreuse-pdt expression plasmid. mChartreuse-pdt expression induced with IPTG (100 μM). mfLon-LOV variant expression induced with arabinose. n = 3 technical replicates.

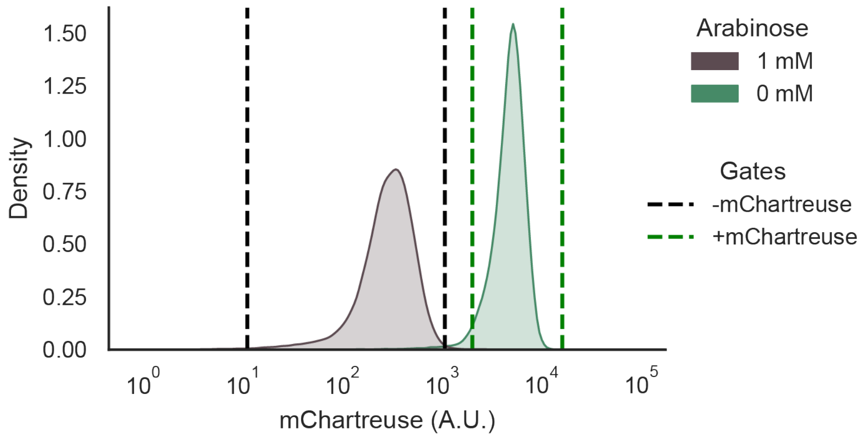


**Figure S9**. Sorting gates were drawn to designate -mChartreuse (black dashed lines) and +mChartreuse (green dashed lines) for enrichment bins. The two populations correspond to mChartreuse-pdt with and without wild-type expression mfLon induced by arabinose.

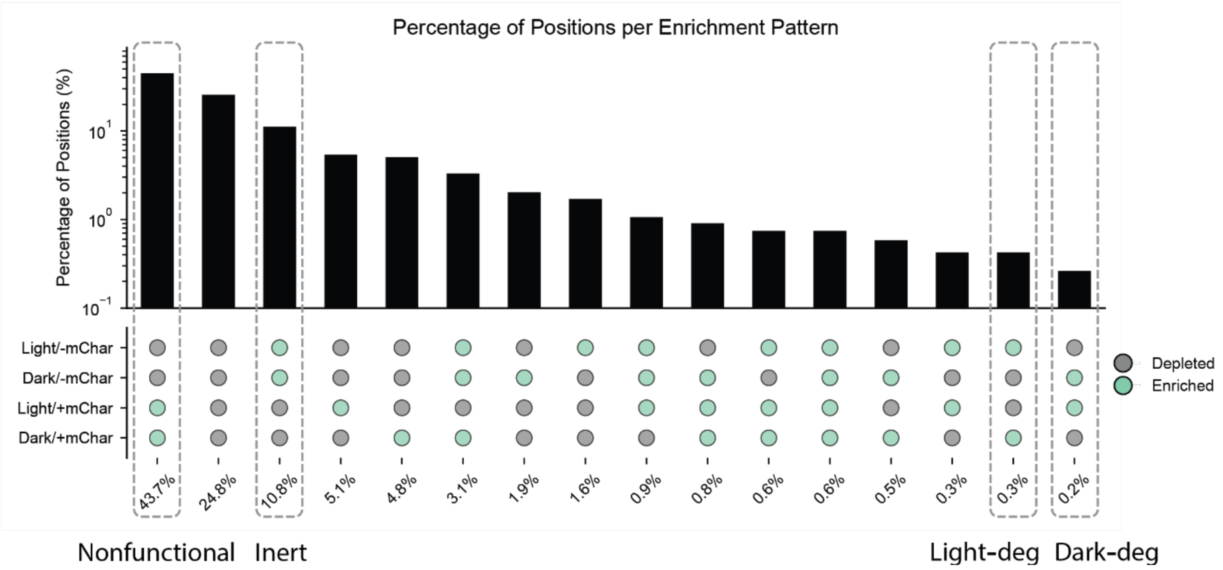


**Figure S10.** Proportion of variants belonging to different combinations of enriched or depleted bins. Enriched corresponds to variants with a positive enrichment score, meaning that, relative to the naïve library, the insertion site is overrepresented in its respective bin. Depleted variants have a negative enrichment score, meaning that, relative to the naïve library, the insertion site is underrepresented in its respective bin. Of the 726 positions, 86 did not appear in the naïve library and were excluded from this distribution. The enrichment combinations that correspond to expected nonfunctional, inert, light-deg, or dark-deg are noted.

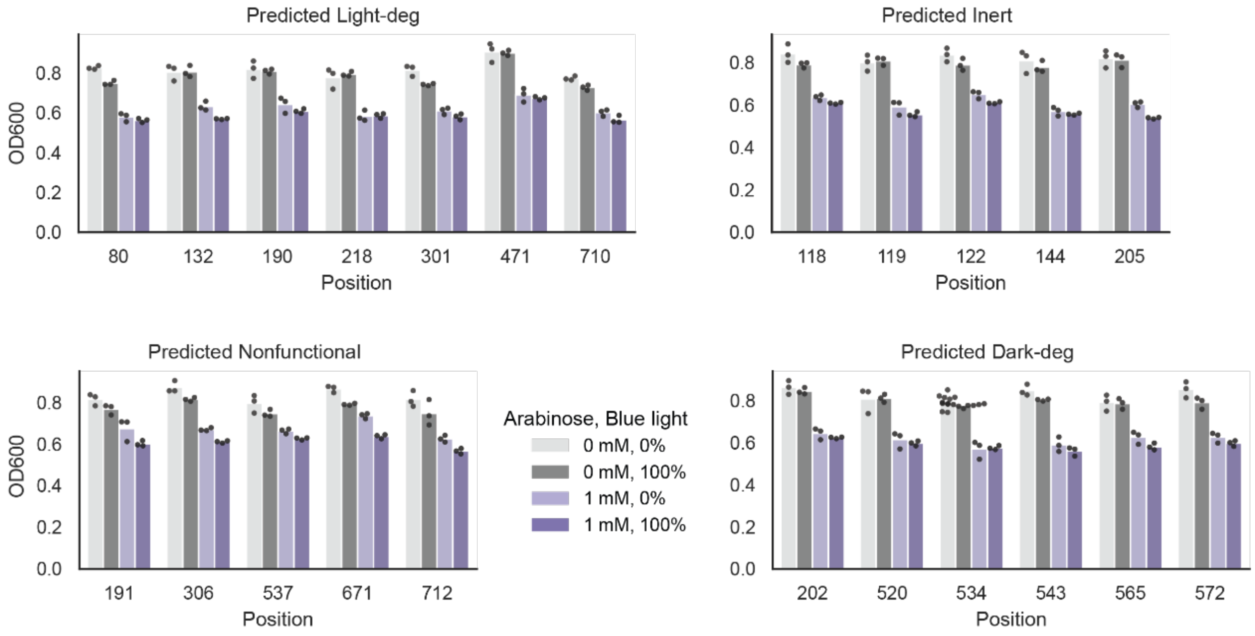


**Figure S11**. Optical density at 600 nm of insertion sites selected through enrichment. mChartreuse-pdt expression induced with IPTG (100 μM). mfLon-LOV-534 expression induced with arabinose. n ≥ 3 biological replicates.

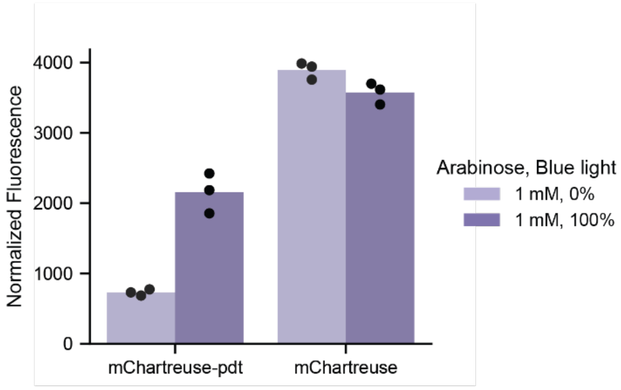


**Figure S12.** Comparing fluorescence readouts from co-transformed strains containing mfLon-LOV-534 and either mChartreuse-pdt or mChartreuse. mChartreuse and mChartreuse-pdt expression induced with IPTG (100 μM). mfLon-LOV-534 expression induced with arabinose. Normalized fluorescence is raw fluorescence divided by optical density at 600nm. n = 3 biological replicates.

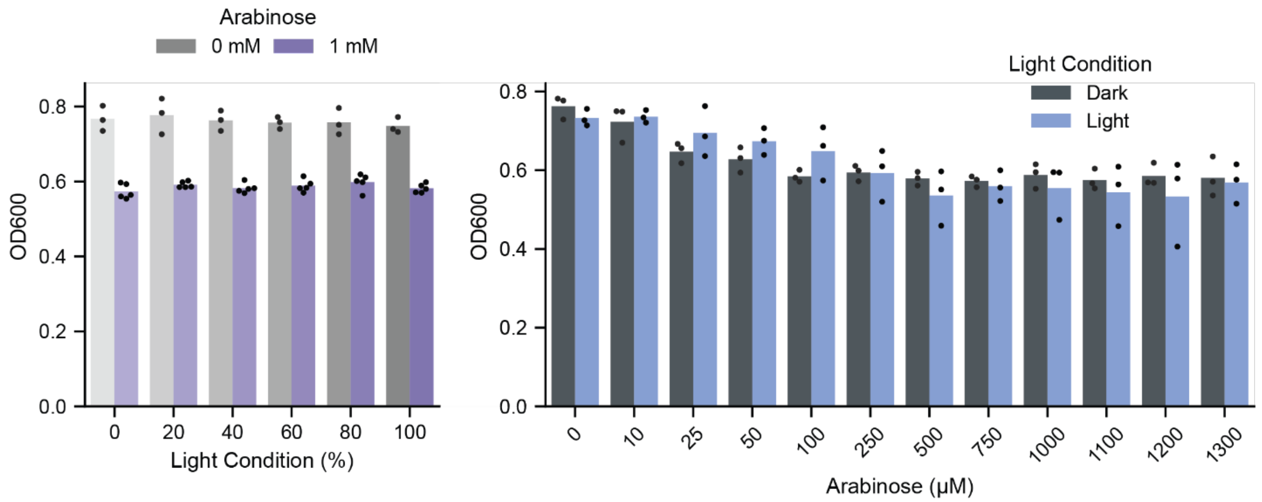


**Figure S13.** Optical density at 600 nm for mfLon-LOV-534 characterization experiments, where we varied light conditions and arabinose. mChartreuse-pdt expression induced with IPTG (100 μM).
